# Long-range actin-driven endosymbiont mobility in a deep-diverging bilaterian

**DOI:** 10.1101/2025.06.08.658386

**Authors:** Grace Zhong, Gaëlle Toullec, Pierre-Henri Jouneau, Johan Decelle, Manu Prakash

**Affiliations:** Bioengineering, Stanford University, 443 Via Ortega, Stanford, 94305, California, United States; Cell and Plant Physiology Laboratory, CNRS, CEA, INRAE, IRIG, Université Grenoble Alpes, Grenoble, 38054, France.; CEA, IRIG-MEM, Université Grenoble Alpes, Grenoble, 38054, France.; Oceans, Stanford University, 443 Via Ortega, Stanford, 94305, California, United States; Biology, Stanford University, 443 Via Ortega, Stanford, 94305, California, United States

**Keywords:** symbiosis, acoel, actin-driven mobility

## Abstract

Symbiosis is everywhere, and “we have never been individuals”[1, 2]. In animal-microbe symbioses, established symbionts are often thought to be confined to a specific cellular or tissue niche[3–7] and generally lose their motile appendages such as flagella[8–15]. However, whether the loss of motile appendages necessarily implies immobility within the animal host remains an open conundrum. Here, we present the discovery of long-range, host actin-driven symbiont mobility in a dinoflagellate-acoel worm symbiosis. Using long-term tracking, fluorescence, and electron microscopy, we find that dinoflagellate symbionts (*Amphidinium sp.*, 10-20***µm*** in size) travel throughout an extensive network of thin host cells (**∼**200 nm in regions without symbionts) in *Waminoa sp.* acoel worms, which are part of a deep-diverging bilaterian lineage[16–18]. Although FIB-SEM-based 3D reconstruction shows symbionts still retain both flagella, we uncover that it is host actin machinery that plays a primary role in overcoming large drag forces under confinement to achieve mobility throughout the worm at surprisingly high velocities (around 1***µm/s***). Long term in-toto imaging further reveals diel rhythms and spatiotemporal regulation of symbionts during regeneration. Our findings show the presence of host-mediated mobility in animal-microbe symbioses, which suggests the existence of previously overlooked regulatory processes in holobionts’ maintenance of dynamic homeostasis.

## 1 Introduction

“Now as throughout Earth’s history, living associations form and dissolve” [1]. In an early formal description of animal-microbe symbioses, Sir Frederick Keeble describes a “plant-animal” - a worm found on the beaches of Brittany that photosynthesizes using green “plant-like cells” [19]. We now know that the “plant-animal” - *Symsagittifera roscoffensis*, endearingly called the Roscoff worm - is an animal and a holobiont. It is an acoel worm - part of a deep-diverging bilaterian lineage[16–18] - that hosts photosynthetic algal symbionts. The field of symbiosis is rapidly evolving, yet one notion remains relatively unchanged: in animal-microbe symbioses, despite the importance of motility in symbiont acquisition[20, 21], established symbionts are considered generally nonmotile[8–14] inside the body of a host. For instance, dinoflagellate symbionts in corals transition from a swimming bi-flagellated form to a coccoid (non-flagellated) form[9, 10, 22–24], which is described as nonmotile[9, 10, 22]. *Vibrio fish-eri* - bacterial symbionts in the Hawaiian bobtail squid - become non-flagellated after acquisition in the squid’s light organ[25] and are thereafter considered nonmotile[11, 12]. Host immune responses in the human gut result in low flagellin expression in commensal gut bacteria, rendering them “generally nonmotile” [13, 14]. However, there is little inquiry into whether lack of symbiont motility apparatus (nonmotile) in animal-microbe symbiosis necessarily implies absence of movement (immobility) within the host, perhaps due to lack of tools for in-toto imaging in most animals.

The surprising lack of knowledge about symbiont mobility in the animal world leaves as open the question as to how highly dynamic holobionts maintain homeostasis in constantly shifting metastable states at organismal, tissue, and cellular scales[26]. For instance, corals have been known to shuffle symbionts in order to survive changing ocean temperatures[27]. These kinds of shifts are hard to explain with immobile symbionts and slow cell division timescales. In single-cells, knowledge of symbiont motility such as in colonial Collodaria[28] opened up new ways of thinking about metabolic interaction between partners and nutritional sources accessible for the symbiont. Knowledge of host-mediated symbiont mobility such as in Foraminifera[29] suggests an important additional level of host regulation in response to factors such as light exposure. Although critical in single cells, symbiont mobility in animal-microbe symbioses has so far been overlooked.

The lack of knowledge on host-mediated symbiont mobility is also surprising given the well-established field of organelle mobility. Both intracellular symbionts and symbiont-derived organelles (mitochondria and plastids) are thought to have lost their motile capacities[5, 8, 13, 14, 23–25]. However, despite being on an evolutionary continuum, organelle mobility is well-established and is controlled by the host cytoskeleton[8, 30–32].

Here we present the discovery of long-range, host actin-mediated mobility of endosymbionts in *Waminoa sp.* (an acoel worm, Fig.1D). Dinoflagellate symbionts, with diameters of 10-20 *µm*, are found to traverse the entire body of the worm, with speeds up to the order of 1*µm/s*. Using in-toto tracking microscopy, we establish the dynamics of symbionts at individual and population scale within the body of a worm. Using 2D Transmission Electron Microscopy (TEM) and 3D Focused Ion Beam - Scanning Electron Microscopy (FIB-SEM), we conclusively establish that symbionts are intracellular, hosted in ultra-thin, sheet-like processes of peripheral parenchyma cells. Symbiotic dinoflagellates travel inside these thin processes forming a network architecture embedded within a unique tissue environment that exists as “negative luminal space” between large, vacuolated parenchymal cells. While dinoflagellates experience a drag force several orders of magnitude higher in these confined thin processes compared to a swimmer in unconstrained cytoplasm, we find that these confinement effects in a host cell also enables actin-based mobility with velocities higher than most other actin-driven systems. Using long term in-toto imaging, we also discover dynamic spatiotemporal distributions of symbionts throughout the animal’s body during diel rhythms and under regeneration. Taken together, our work establishes a unique mechanism for symbiont mobility inside the complex tissue architecture of an acoel worm, an important milestone in animal evolution.

**Fig. 1.**
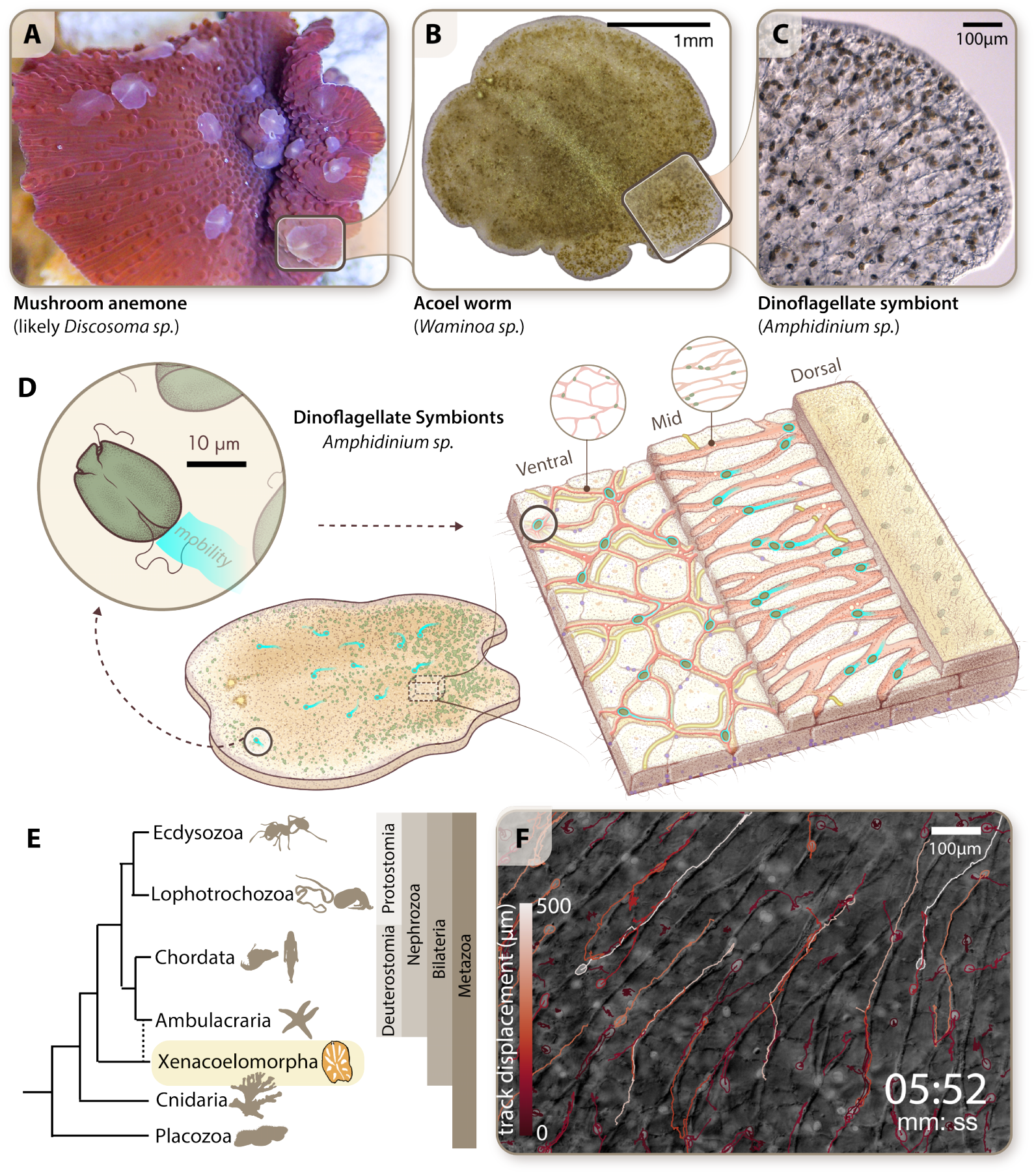
Symbiont mobility in *Waminoa sp.*, a deep-diverging bilaterian. **(A-C)** The system we study is part of a “three-party symbiosis”[42]. (A)*Waminoa sp.* preferentially rests on corals and mushroom anemones. They capture live plankton for food and are hypothesized to consume mucus from host corals [42, 47]. (B) *Waminoa sp.* hosts dinoflagellate symbionts (*Amphidinium sp.*) which are spread all throughout the worm. (C) Symbionts are 10-20 *µm* in diameter. **(D)** Acoel worms have simple body plans characterized by the lack of a through-gut and a coelom [17, 34], and tissue architectures are highly diverse within acoelomorpha [34–41]. *Waminoa sp.* has a ciliated dorsal and ventral epithelium. The epithelium comprises of in-sunk cells, with a layer of neurons and body wall muscles just beneath. The middle of the worm is largely occupied by large central parenchyma cells with giant vacuoles, which are fenestrated by peripheral parenchyma cells (with long and thin processes forming a spanning network throughout the animal) and sparse host musculature. Via live microscopy, we establish two key layers of interest in this study: termed “ventral” and “mid” layers. The ventral layer lies right above the epithelium and muscle layer. In this layer, peripheral parenchyma cells exist alongside many other host cells, such as muscles, neurons, in-sunk nuclei of epithelial cells, and neoblasts. The layer contains many host nuclei. The mid layer is 30-60*µm* inwards from the ventral layer, depending on the thickness of the worm. It is morphologically distinct from the ventral layer in that it is composed mostly of large, vacuolated central parenchyma cells. Peripheral parenchyma cells, dinoflagellate symbionts, and sparse host musculature are relegated to the negative space between these large cells. **(E)** Acoel worms are likely extant members of the earliest-diverging bilaterians, sister to nephrozoa (all other bilaterians)[16, 18]. An alternative hypothesis places them as sister to ambulacraria[17]. In either case, acoel worms have been recognized as a key evolutionary group in context of body plan development in metazoans. **(F)** We discover long-range directed mobility in dinoflagellate symbionts in *Waminoa sp.*. Symbionts travel along “transport highways” visible under DIC microscopy. “Highways” can accommodate multiple symbionts and support bidirectional travel. There are often “lane switchings”, fast turns, collisions, and passing events (SI Video 1). The longest trajectory of a dinoflagellate we are able to track for the entire duration is 700 *µm* over 6 minutes (white tracks, SI Video 2). This is an underestimate of the maximum distance traveled since all dinoflagellates continue to move in the bulk tissue. Thus, the above metric represents the limit in our ability for long term tracking a single cell – dinoflagellate live inside a healthy motile worm (See Methods).

## 2 Results

### 2.1 Acoel worms as a model system for studying symbiosis

Acoel worms occupy a unique phylogenetic space as part of Xenacoelomorpha - a deep-diverging lineage[33] which is thought to be either extant members of the earliest diverging bilaterians[16, 18] or sister to Ambulacraria [17] (Fig.1E). Their simple body plan lacks a through-gut and a coelom (Fig.1D) [17, 34]. Importantly, tissue architectures are highly diverse within acoelomorpha [34–41], which highlights the clade’s unique phylogenetic position for understanding the evolution of cell types and tissue architecture[34], and for investigating the role of symbiosis in shaping these processes.

The holobiont (*Waminoa sp.*) that is the focus of our study is part of a unique “three-party symbiosis” between a coral, the acoel worm *Waminoa sp.*, and dinoflagellates (SI Video 2, Fig.1A-C)[42]. *Waminoa sp.* has been reported to associate with many host coral species in reefs of Singapore[43], Australia[44], Taiwan[45], the Red Sea[42, 46], Japan[47], and Indonesia[48]. While the worms live on coral surfaces, living inside the tissue of these worms are dinoflagellates of the genus *Amphidinium*[49], with the co-presence of *Symbiodinium* species distinct from the coral symbiont reported in several studies[42, 47]. The specific acoel worms collected for our study were found atop the mushroom anemone *Discosoma sp.* (see Methods for details) and contain only *Amphidinium sp.* symbionts. We establish stable cultures of *Waminoa sp.* in the lab to enable the study of the acoel worm-dinoflagellate symbiosis and focus on the cellular scale dynamics of symbionts across the entire animal host (see Methods).

### 2.2 Dinoflagellate symbionts of *Waminoa sp.* demonstrate long-range directed mobility

We performed long term, tracking microscopy in live *Waminoa sp.* worms at room temperature and under standard white light illumination (more than 20 individuals, see Methods for details) using SQUID imaging platform capable of tracking a single mobile cell deep inside the tissue of a live animal[50]. In this unique frame of reference of individual worms, we observe long-range mobility of symbiotic dinoflagellates inside the host tissue (SI Video 1, Fig.1F) throughout the animal. Specifically, the symbionts - 10-20*µm* in diameter - are capable of moving hundreds of micrometers in distance over the timescale of minutes with ballistic dynamics while the host remains completely still (Fig.1F, SI Video 1). Moreover, no detectable correlation is observed in symbiont and worm mobility. Dinoflagellate trajectories also overlap with vesicle “transport highways” visible under differential interference contrast (DIC) imaging (Fig.1F, SI Video 1). Evident in SI Video 1, the dynamics of symbiont mobility are highly diverse. We observe bi-directional mobility along a network of transport “highways”. Cargoes are seen to switch “highways” at junctions, hinting at the presence of a syncytium. Surprisingly, symbionts also switch directions rapidly. Large dinoflagellates are seen to cross and pass each other with natural ease. We also observe rapid overtaking events, as well as direct cell-cell collisions. Collisions can also result in directional reversal. Qualitative observations also reveal reduced mobility in the animal’s periphery as compared to bulk tissue, creating an analogy to traffic jams occurring at cellular scale.

Via high-resolution tracking microscopy, the longest single cell displacement inside the host tissue we capture in our experiments is 700 *µm* over a period of six minutes (Fig.1F, SI Video 2). This is an underestimate of the maximum distance a dinoflagellate can travel in the host because it reflects a limit in our ability to track the symbiont’s 3D movements within the host’s tissue (See Methods). We also record single dinoflagellates switching directions rapidly, without any prolonged deceleration or ensuing acceleration (Fig.2E-F, SI Video 2), a common hallmark of low Reynolds number hydrodynamics. Interestingly, when dinoflagellates change direction of motion, we do not observe change in heading (rotation of the cell), implying the mobility in this case is independent of cell polarity. This long-range mobility stands as an outlier in previously known mechanisms for symbiont assimilation throughout animal host tissue. We next seek to determine the tissue niche in acoel worms that hosts these mobile symbionts, and if the mobility is observed across the multi-layer tissue architecture of the worm.

**Fig. 2.**
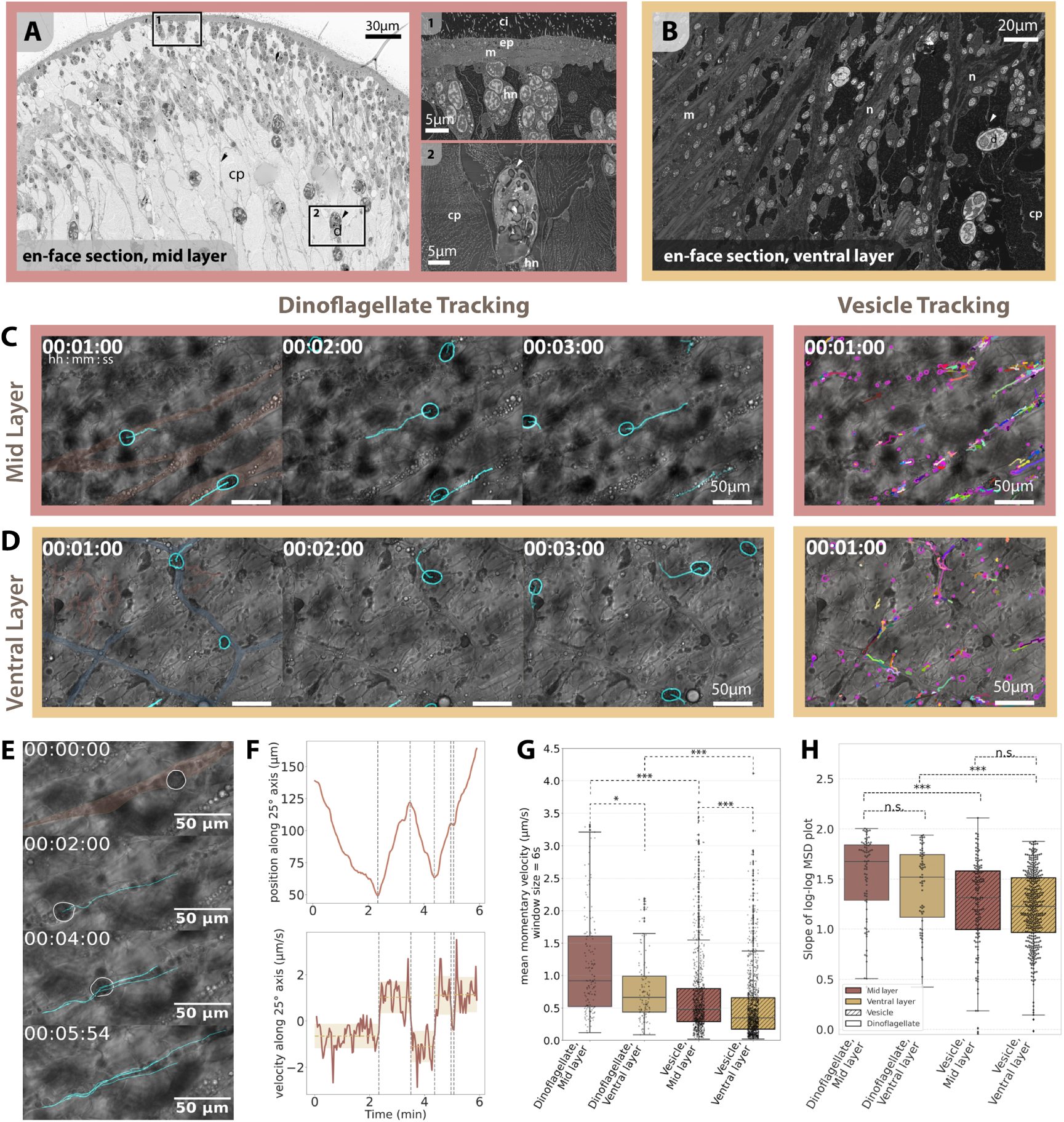
Symbiont mobility occurs in morphologically distinct layers of the worm. (**A-B**) En-face scanning electron microscopy (SEM) sections of the worm’s mid (A) and ventral layers (B) highlight similarities and differences between the two layers (represented in Fig.1D). In both layers, dinoflagellates are found inside the thin processes of peripheral parenchyma cells (arrowheads). Host nuclei for these parenchymal cells as well as for other cells are enriched at the worm’s periphery (A, inset 1), with abundant host nuclei in the ventral layer (B). The ventral layer is also traversed by host neurons, whereas the mid layer is characterized by higher numbers of symbionts and sparse host musculature relegated to the “negative spaces” between central parenchyma cells containing large inflated vacuoles. ep: epithelium, m: muscle, ci: cilia; n: host neurons, cp: central parenchyma; d: dinoflagellate; hn: host nucleus (**C-D**) In both mid (C) and ventral layers (D), DIC tracking microscopy reveals that dinoflagellates and vesicles share the same “transport highways” (examples highlighted in maroon in the first frame of (C) and (D)). “Transport highways” are easy to discern and wide in the mid layer (C) and harder to discern in the ventral layer (D). In the ventral layer, the directionality of dinoflagellates appears to be somewhat correlated with the polygonal-shaped host neurons (highlighted in blue in the first frame, SI Fig 1). See SI Video 3. (**E**) Via tracking microscopy, we show a sample trajectory of a dinoflagellate in the mid layer (“transport highway” highlighted in maroon). The dinoflagellate reverses directions three times in this dataset over a period of six minutes. (**F**) Surprisingly, direction switching occurs without changes in heading (SI Video 2) and without prolonged deceleration or acceleration. Regardless of direction of motion, the mean momentary velocity in each segment of motion is similar at around 1*µm/s*. (**G**) By comparing datasets from individual dinoflagellates in the mid and ventral layers, we find that symbionts move significantly faster in the mid layer as compared to their counterparts in the ventral layer. Although dinoflagellates are significantly larger than individual vesicles, they still move significantly faster than vesicles in both the layers. (**H**) Measuring directedness of mobility using slopes of log-log plots of mean squared displacement (MSD) vs time interval, we find that dinoflagellates in both the mid and ventral layers show nearly ballistic motion, typical of active driven cellular processes. In both layers, dinoflagellate motion is significantly more persistent than that of vesicles. ∗ : 0.01 *< p <* 0.05, ∗∗ : 0.001 *< p <* 0.01, ∗ ∗ ∗ : *p <* 0.001

### 2.3 Directed symbiont mobility is pervasive throughout host tissue

In order to address whether symbiont mobility occurs throughout the tissue of the host, we first highlight unique aspects of worm tissue architecture in *Waminoa sp.*. Acoel worm tissue architecture has been previously described qualitatively[35, 46, 51–54]. As illustrated in Fig.1D, in cross section, *Waminoa sp.* has multi-ciliated dorsal and ventral epithelium layers. It lacks a coelom and a through gut, and only a single orifice for feeding is found on the ventral surface. Inwards from the epithelium layer, there are neuronal and body wall muscle layers[51]. Further inwards, acoel worms are generally understood to have an ill-defined central parenchyma, penetrated by long processes of peripheral parenchyma cells[52, 53]. Some acoel worms contain symbionts, which are typically located in this parenchyma[46, 54]. To better map this unique high-density cellular architecture, we employ TEM and FIB-SEM to both corroborate previous descriptions of tissue architecture and map symbiont localization across the animal host.

Acoel worm geometry can be understood as a flat pancake-like architecture with multiple layers. In order to situate our data in these morphologically distinct layers within the acoel worm’s parenchyma, we focus on two key layers: ventral layer and mid layer (Fig.1D). The layers can be easily seen both via DIC imaging in live animals and also in TEM and FIB-SEM reconstructions. The ventral layer lies just above the ventral epithelium and muscle layer. Here, peripheral parenchyma cells exist alongside many other host cells, such as muscles, neurons, in-sunk nuclei of epithelial cells, and neoblasts (Fig.1D, Fig.2B). The mid layer (Fig.1D, Fig.2A) is found 30-60*µm* further inwards from the ventral layer, depending on the thickness of the worm. This layer contains fewer host nuclei save for at the worm’s periphery. Our TEM datasets (Fig.3A-E) establish that it is composed mostly of large, vacuolated central parenchyma cells (highlighted in purple in Fig.3A-E). Remarkably, we discover that the thin processes of peripheral parenchyma cells and musculature in this layer are almost pinned to the negative luminal space between large vacuolated cells which take up the majority of the space - akin to negative space between balls in a packed ball-pit (Fig.3A-E). This fenestrated co-joint morphology between vacuolated cells and thin peripheral parenchymal cells can be understood as a linked topological architecture, previously also reported at a cellular scale in ultra-large single cells[55].

**Fig. 3.**
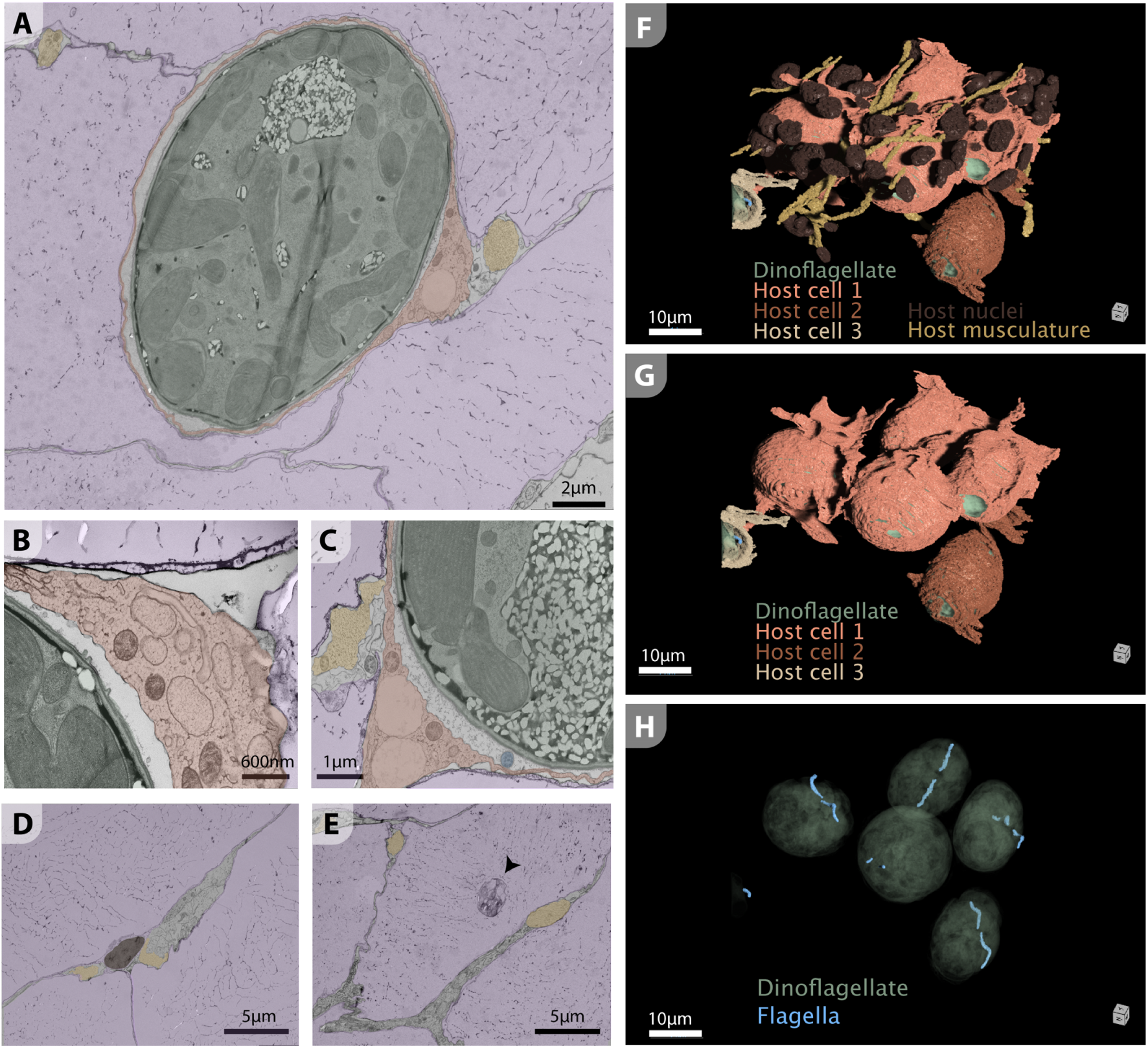
Intracellular nature of symbionts within host cells. **(A-E)** TEM images preserving the delicate cellular membranes reveal that symbionts (green) are completely inside host peripheral parenchymal cells (pink). The host cells with thin processes forming a network exist within the negative space of large, inflated, and highly vacuolated central parenchymal cells (purple), along with sparse worm musculature (yellow). For several datasets, the symbiosome is captured as the space between the symbionts (green) and the host cells (pink) (A-C). We also observe unidentified filamentous materials inside the symbiosome (more examples in SI Fig.3). Host nuclei are sparse (brown, in D) and may belong to a large vacuolated cell. In some large vacuoles, we see what could be lysosomal vesicles (arrowhead in E), suggesting that these giant-vacuole-containing cells (purple) may be part of the central digestive parenchyma. To establish the true localization of symbionts within the host cells, we perform 3D volumetric reconstruction of smaller tissue sections via FIB-SEM (see Methods). **(F-G)** Segmented 3D FIB-SEM volume confirms the intracellular location of symbionts and reveals that the cells hosting symbionts have thin, sheet-like, processes, matching previous descriptions of a subset of peripheral parenchyma cells [35, 56] and what we observe in live movies of symbiont mobility presented earlier (SI Videos 1-3). Multiple symbionts can reside in a single host cell, akin to traffic jams (at least four symbionts depicted within a single host cell). The host cells we delineate extend beyond our volume. Though there are many host nuclei in the ventral layer (F), our volume does not contain the nuclei of any of the three cells hosting symbionts. **(H)** Surprisingly, all five dinoflagellates that are fully contained in the segmented FIB-SEM volume clearly retain both flagella. Although flagellar length varies across different symbionts, both flagella are still positioned correctly in the corresponding grooves in the theca.

Via live imaging of worms, we further establish that the network architectures of transport highways in the ventral and mid layers are geometrically distinct (Fig.2A-B). Comparing DIC videos of symbionts moving in the two layers (SI Video 3), in the mid layer, “highways” appear wider and more persistent (Fig.2C). In the ventral layer, “highways” appear thinner and harder to discern, co-existing with another polygonal network that guides the dinoflagellate trajectories (Fig.2D). Using immunostaining, we find these polygonal paths to be host neurons (SI Fig.1). Further quantifying symbiont mobility in the two layers reveals that both dinoflagellates and passive vesicles achieve significantly higher mean momentary speed (see Methods for details) in the mid layer as compared to the ventral layer (Fig.2G). Although dinoflagellates are much larger in size with significantly more hydrodynamic drag when compared to the small vesicles being shuttled, we find that these symbionts move significantly faster than vesicles in both layers (Fig.2G).

Moreover, dinoflagellate mobility is nearly ballistic at the timescale considered (minutes) in the highly confined mid and ventral layers (Fig.2H) almost resembling 1D microfluidic channels (highways). In both layers, however, we find that the slope of the log-log mean-squared-displacement (MSD) versus time is significantly lower for vesicles as compared to dinoflagellates. Thus, vesicle motion in these highways is less directed than dinoflagellate mobility. To understand the fundamental mechanism for symbiont mobility, we next look at what constitutes these transport networks and the exact nature of symbiont micro-environments within the animal.

### 2.4 Symbionts are intracellular, traveling inside host peripheral parenchyma cells

In order to truly establish the physical environment of these cellular cargo trains, it is crucial to map and reconstruct the 3D tissue architecture at sub-cellular scale. Thus we turn to volumetric reconstruction of serial FIB-SEM sections in pressure-frozen worms. We first seek to definitively establish whether symbionts in *Waminoa sp.* are intra- or extra-cellular. Previous studies have remained inconclusive on this question [37, 42, 46, 49, 54] due to lack of 3D imaging tools at the nanoscopic scale. Using cryogenic sample preparation (high pressure freezing of live worms and freeze substitution), followed by TEM and FIB-SEM (See Methods for details), we conclusively establish that symbionts of *Waminoa sp.* are intracellular, encased within a symbiosome membrane (Fig.3A-C, F-G). The symbiosome has a distinct volume surrounding the symbionts residing completely inside the host cell (Fig.3A-C). The symbiosome also contains an unknown filamentous material in several occasions (Fig.3C, more examples in SI Fig.3). Analyzing the 3D volume reconstruction close to the periphery of the worm, we catch a cellular traffic jam of dinoflagellates (Fig.3F-H), where four symbionts are seen in close proximity within the same host cell (Fig.3F-G). Host cell processes resemble thin sheets (Fig.3F-G). This further supports the hypothesis that cells housing dinoflagellates are indeed a previously described subtype of peripheral parenchymal cells in acoels characterized by long, sheet-like processes[35, 56]. In these nanoscale 3D high-resolution reconstructions, complete cells hosting symbionts cannot be captured. We were also unable to capture any nuclei of the three segmented host cells.

### 2.5 Symbionts retain both flagella, but flagella do not drive symbiont mobility

We next ask whether it is the host or the dinoflagellates - or both - that control the observed surprising symbiont mobility in our worms. Analyzing our FIB-SEM volume data for flagella, we find that all segmented dinoflagellates retain both flagella - some longer than others (colored in blue in Fig.3H). This is in contrast to prior reports where symbionts are typically known to lose their flagella upon establishment of symbiosis. Within acoel worms, for example, *Tetraselmis sp.* symbionts in *C. longifissura* and *S. Roscoffensis*[57] lose all four flagella in their relationship with their respective host acoel worms [58, 59]. There is only one past report on retention of both flagella in an acoel symbiont, which is in the association between the dinoflagellate *Amphidinium klebsii* and the acoel worm *Amphiscolops langerhansi* [60]. In this particular host-symbiont system, the symbiont is believed to be extracellular, and their extracellular nature is hypothesized to be an explanation for their flagella retention [60].

Given the endosymbionts’ retention of flagella, we next explore whether the observed mobility could be driven by symbiont flagella. First, in single cell tracking datasets, we do not observe the classical on-axis rotation of a swimmer being propelled by a flagella (represented as a stokeslet in low Reynolds number regime). The rotation of a cell during a cycle of flagellar stroke is known across the eukaryotic tree of life and is also characteristic of free swimming dinoflagellates driven by flagellar dynamics [61]. Strong evidence of mobility without cellular rotation is visible in “photo-marked” dinoflagellate cells in confocal live imaging datasets (SI Video 5) where autofluorescence marks on symbionts (Fig.4F) traverse the cell without any apparent rotation. Secondly, we do not observe any changes in cell orientation when the dinoflagellate suddenly changes directions (SI Video 2). Although many flagellates are known to show a certain degree of reversibility while swimming, they often turn and utilize the propulsive flagellar movement, favoring a single polarity. In high-magnification live imaging videos, we also fail to observe any flagellar beating, which is necessary for flagella-driven motility. Correlating our live datasets with our 3D volume reconstruction, we note that all dinoflagellates are found within an internal membrane forming a true symbiosome encapsulated inside the host cells. This generates a topological barrier where any potential flagellar beat generated inside a symbiosome within this tight confinement of another cell cannot enable momentum transfer to the cytoplasm of the host cell (outside the symbiosome), which would be essential for motility relative to the host cell[62]. This suggests flagella are not responsible for the observed symbiont mobility. We next look for host cell-based mechanisms.

**Fig. 4.**
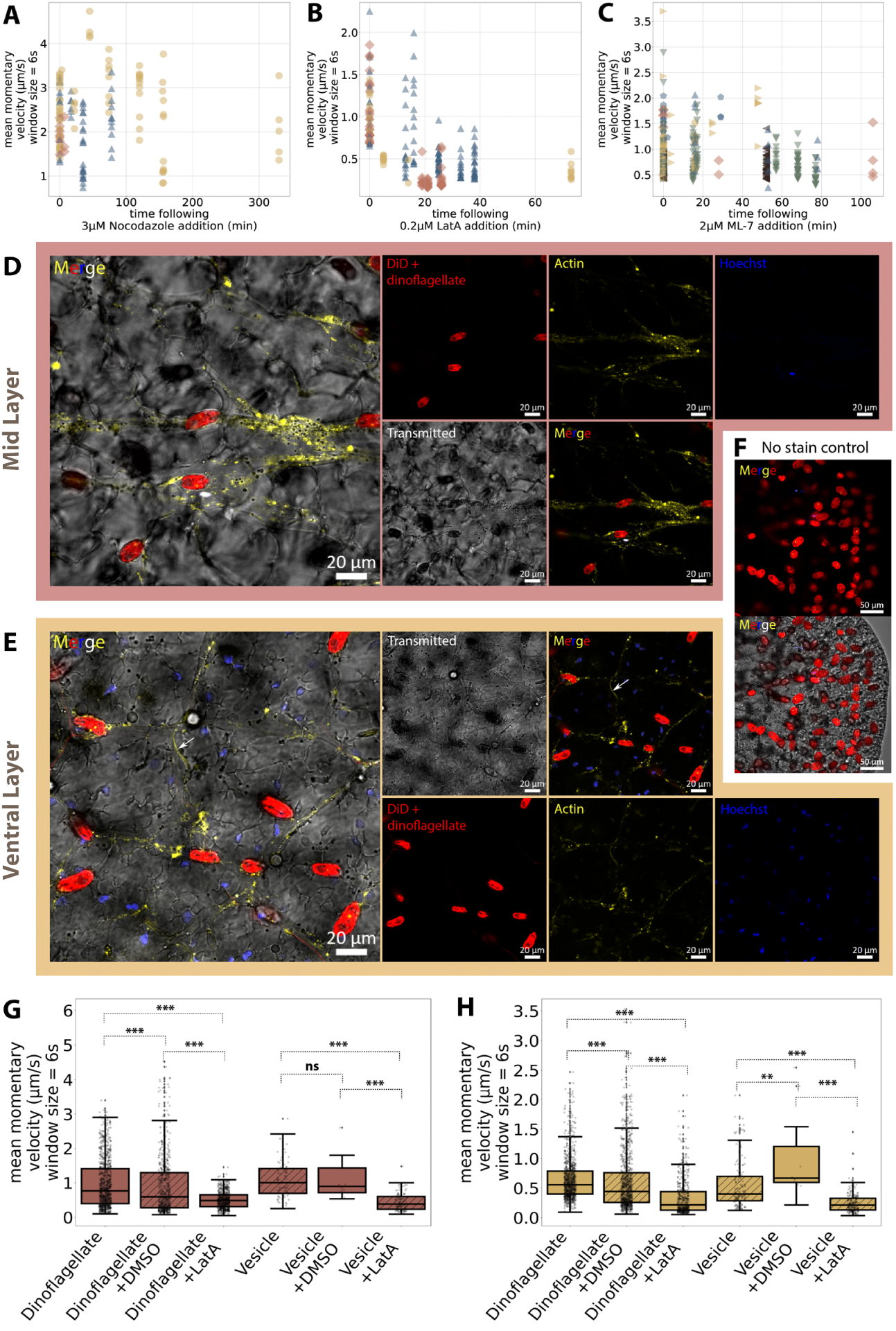
Host actin plays a key role in symbiont mobility. To elucidate the role of host cytoskeleton, we explore drug perturbation assays to modulate symbiont mobility in living organisms. **(A)** We do not find any significant inhibition of symbiont mobility with the addition of 3*µM* nocodazole (a microtubule disrupting drug[63]), suggesting that the mobility is not microtubule-driven. **(B)** Addition of 0.2*µM* LatA (common drug which inhibits actin polymerization by sequestering G-actin[64]) is enough to cause fast and marked inhibition of symbiont mobility (SI Video 6), suggesting a key role of actin. **(C)** Addition of 2*µM* ML-7 (a specific inhibitor of myosin light chain kinase[65, 66]) also inhibits symbiont mobility, albeit to a lesser degree than 0.2*µM* LatA, suggesting potential yet limited role of myosin (SI Video 6). Each marker type in A-C represents data from a different worm. **(D-E)** In order to truly elucidate the molecular underpinning of this mobility, we turn to live confocal imaging with live actin stain (CellMask^TM^ Orange Actin Tracking Stain). Surprisingly, live confocal microscopy reveals a large amount of F-actin present in host cells that harbor symbionts, in both the mid (D) and ventral (E) layers. We can clearly visualize actin puncta of various sizes traversing and flowing through the exact same transport highways that host the symbiont cells. As shown in SI Video 5, these puncta flow bidirectionally in the cytoplasm of host cells containing symbionts, changing directions as do both vesicles and symbionts. Although these actin puncta are highly enriched in these specific symbiont-containing host cells, they are hardly present anywhere outside the specific “transport highways”, suggesting actin filaments within the host cytoplasm play a role in symbiont mobility. Actin bundles (arrow in merged images, E) are also seen on the ventral layer in other (likely muscle) cells where they are static (SI Video 5). Hoechst-stained nuclei in both the layers do not appear to move (SI Video 5). **(F)** Dinoflagellate rotation can be further tracked qualitatively in these datasets due to the presence of a TRITC autofluorescence “dot” (yellow) on the symbiont. We observe no rotation during mobility across the host cell (SI Video 5). **(G-H)** Addition of 10*µM* LatA significantly inhibits dinoflagellate and vesicle mobility in both mid (G) and ventral (H) layers (SI Video 6). DMSO used as a control has a minor effect on mobility, which is significantly smaller when compared to LatA. ∗ : 0.01 *< p <* 0.05, ∗∗ : 0.001 *< p <* 0.01, ∗ ∗ ∗ : *p <* 0.001

### 2.6 Host actin is the primary driver of symbiont mobility

Since the system under study is not amenable to genetic perturbations, we perform live imaging and drug perturbations to explore the role of host cytoskeleton in symbiont mobility. Host cytoskeletal components, namely microtubules and actin, often play major roles in organelle mobility [67, 68]. Long distance transport inside cells such as neurons is primarily driven by microtubule-based motor transport[69], though recent work also highlights the role of microtubule- and actin-based mobility of host nuclei (albeit slow) in neurons [70, 71]. These examples highlight that mobility can be achieved under confinement, a hallmark of thin peripheral parenchymal cells we focus on in our study. Inspired by this, we treat the animal with 3*µM* nocodazole (a microtubule disrupting drug[63], see Methods for details). Nocodazole treatment has no discernible effect on symbiont velocity over five hours (Fig.4A), ruling out the role of host microtubules.

We next focus on actin and its role in symbiont mobility within the worm. More specifically, we test two key hypotheses comparing actin polymerization-based mobility (often referred to as actin comet tails) [70, 72–75] versus actomyosin-based mobility[76]. First, we use live actin imaging under confocal microscopy to observe actin dynamics in these unique ribbon-like cells that host symbionts. We choose CellMask Orange Actin Tracking Stain (Fisher Scientific, A57244), which provides strong fluorescent labeling of polymerized/filamentous actin in live organisms and is well-tolerated by our worms (no physiological changes observed over days). In the ventral layer (SI Video 5, Fig.4E), both static actin bundles (from host musculature) and flowing actin puncta (inside host cell cytoplasm) are clearly visible. Transport highways as shaped by flowing actin puncta are weakly correlated with, but separate from, thin processes labeled by DiD (membrane and lipid stain, Fisher Scientific V22887), which are possibly neuronal processes across the ventral layer (SI Fig.1). Many host nuclei are also visible in this layer. In the mid layer (SI Video 5, Fig.4D), the actin puncta appear far denser, and the “highways” as clearly outlined by flowing puncta appear thicker and throughout the host cell network. This also correlates with higher symbiont velocities in the mid layer (Fig.2G), suggesting a role of host actin in symbiont mobility. There are noticeably very little host nuclei. DiD does not stain any cells in this layer.

Focusing on live actin flow in host cells, we find that both layers exhibit actin puncta of various sizes moving coherently as a collective. Although hard to track automatically, we manually measure and estimate the velocity of the actin puncta to be roughly 1.0 ± 0.7*µm/s* across six puncta analyzed. Small vesicles or droplets are also seen zipping with actin puncta (panels merged with transmitted images in SI Video 5 and Fig.4D-E). We observe bi-directional fluxes and junctions at which labeled actin puncta - along with their cargo - can “make turns”, “switch lanes”, or bypass one another. The dynamics of actin puncta is indeed similar to dinoflagellate mobility inside these same host cells.

To better understand how actin drives mobility, we next perturb this robust mobility phenotype using drugs targeting actin-polymerization based mobility. Latrunculin A (LatA) is a common drug that has been used extensively to inhibit actin polymerization by sequestering G-actin[64]. When 0.2*µM* LatA is added, while a few small vesicles continue to move, almost all dinoflagellate motion is clearly and suddenly halted (SI Video 6). Adding a higher concentration of 10*µM* LatA and observing using live actin stain under confocal microscopy, we further observe remarkable cellular traffic jams where LatA addition clearly stalls both the flow of actin puncta and the associated symbionts (SI Video 6). At this higher concentration, we also observe a unique phenotype of dinoflagellate clumping (SI Video 6), possibly due to traffic jams and passive minimization of membrane deformations in a clustered state of dinoflagellates inside tubular membrane processes of the host cells.

We can further quantify this striking visual difference in mobility (Fig.4G-H). We find that 10*µM* LatA addition causes a significant decline in velocity in both mid and ventral layers for both dinoflagellates and host vesicles. At a lower concentration of 0.2*µM* LatA, we still observe a fast and sharp inhibition of symbiont mobility within a mere tens of minutes after LatA is added (Fig.4B). The remarkably sharp inhibitory effect of even a low concentration of LatA strongly suggests that host actin polymerization plays a key role in the observed symbiont mobility.

Interestingly, addition of CK-666, which suppresses branched actin network formation by inhibiting actin-related protein 2/3 (Arp2/3)[77], qualitatively leads to similar phenotypes as LatA. Specifically, we observe CK-666 addition to halt symbiont mobility at low concentration (10*µM*) and cause a similar symbiont clumping phenotype at high concentration (50*µM*) (SI Video 6). Unfortunately, CK-666 addition also induces the worm to be very active, so we were not able to track single dinoflagellates inside the body of a live worm to quantify these effects.

To establish if actomyosin-based mobility might also be at play in our system, we use ML-7, a specific inhibitor of myosin light chain kinase[65, 66], and look at symbiont mobility. We find 2*µM* ML-7 does slow down dinoflagellate mobility, and this inhibition is weaker and less consistent across worms (Fig.4C, SI Video 6). ML-7 addition also produces an interesting phenotype - different from LatA - on the timescale of hours where the large vacuolated cells appear more rounded, giving the animal’s tissue a “foam-like morphology” (SI video 6). Addition of 10*µM* blebbistatin - a myosin II inhibitor[78] - has no discernible effect on symbiont mobility (SI Fig.2).

Taken together, our drug perturbation assays and live actin imaging conclusively establish that host actin plays an integral role in the observed symbiont mobility. The visual similarity between the CK-666 and LatA phenotypes suggests the possibility of mobility driven by pushing forces generated by formation of actin networks linked to the symbiosome, perhaps akin to actin comet tails in *Listeria* motility[72–75, 79]. What differentiates these actin tails is the fact that mobility occurs in our system under an extremely high confinement within thin ribbon-like host cells. The weaker and visually different ML-7 phenotype, the lack of effect of blebbistatin, as well as our observation of flowing puncta, suggest that although myosin may be involved in mobility, the main mechanism is not actomyosin-based contractility, and the likely force generation mechanism is actin polymerization. The bidirectional flows and multitudes of possibilities at turns and during cargo collisions, as seen in live DIC movies, can also be explained by actin polymerization linked to the symbiosome surface. Further, the apparent independence of mobility directions of individual symbionts in close proximity (e.g. passing each other in opposite directions) can be explained by a model where actin polymerization provides a propulsive force, with a nucleation site on the symbiont (cargo) determining the final direction of movement. The proposed actin polymerization mechanism is also consistent with the fact that the mobility seen is highly ballistic. Next we explore if it is physically feasible for actin polymerization to drive mobility at the speeds we observe under a high degree of confinement.

### 2.7 Role of confinement in symbiont mobility

In the wide array of imaging modalities presented in our work including low and high magnification videos (SI Videos 1-3) and FIB-SEM-based reconstruction, it is clear that symbionts are rapidly moving inside thin processes of host cells under a surprisingly high degree of confinement. In terms of geometry, encapsulated dinoflagellates in *Waminoa sp.* would experience extreme degrees of confinement inside these ribbon-like host cells (Fig.3F-G) which are only 35-200nm in width in areas without symbionts([56], also measured from our FIB-SEM dataset). Yet we find that host cells are able to stretch rapidly to accommodate traveling dinoflagellate cargoes as large as 10-20 *µm* in diameter (nearly two orders of magnitude larger than itself). We explore what effective drag forces might be present in such high degree of confinement and how the system is able to generate high propulsive forces for this remarkable cellular mobility inside a living host.

Two relevant comparative examples of transport under high degrees of confinement are nuclear migration inside neurons[70, 71] and organelle transport in tunneling nanotubes (TnTs)[80–82]. In both examples, both actin and microtubules have been implicated[70, 71, 80, 82, 83]. In the case of nuclear migration inside neurons, axonal processes range from 1.5-3*µm* wide with axial symmetry, and nuclei are about 5-10*µm* [70, 71]. As compared to dinoflagellates with stiff thecal plates, nuclei are far more deformable. Past work on nuclear migration within neurons finds traversal velocities of only 17nm/s[70, 71], significantly lower than what we report in our work. In the second example of mobility under confinement, we look at the case of organelle transport within much smaller membrane nanotubes (TnTs) with tube diameters of 50-1000 nm [80–82]. Typical organelle cargo such as mitochondria are about 11*µm* in diameter[84], and cargo velocity has been typically reported to be only 25-100nm/s [80, 83, 85]. In one study of quantum dot transport in TnTs, velocities similar to what we report in our study (1.23*µm/s* [82]) have indeed been reported, though quantum dots are significantly smaller and hence do not experience the same degree of confinement as in our current study.

Another comparison to keep in mind is the example of Arp2/3 mediated, actin-polymerization-driven transport in generating comet tails that propel a cell (*L. monocytogenes*) forward inside the cytoplasm of another cell [72–75]. Although this example does not have the same degree of confinement as presented in ultra thin processes of peripheral parenchymal cells in our study, it is still valuable to compare velocities previously reported in actin comet tails, which is roughly 0.1*µm/s* - an order of magnitude lower than what we find (1*µm/s*) in *Waminoa sp.*. These comparisons indeed present an interesting conundrum, where we find a much faster mobility of large symbiont cells in such a high degree of confinement, which presents a higher drag (resistance to movement).

Indeed, a past theoretical model exploring cargo transport in narrow tubular membrane structures such as TnTs predicts a rapid increase in drag force associated with increasing cargo size under high confinement[86]. For membrane nanotubes of rigidity *k* and surface tension *σ*, free energy of a nanotube comprises of bending energy 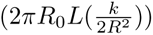 and surface tension (2*πR*_0_*Lσ*). At equilibrium, balancing these two forces provides an average membrane tube radius 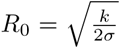. For typical membrane bending energy of *k* = 4.10^−20^*J* and *σ* = 2.10^−6^*Nm*^−1^, membrane tubes can easily be as thin as *R*_0_ = 100*nm*[87]. In order to compute drag of a particle in such a nanotube, several factors have to be included. Drag forces faced by a cargo traveling inside a complex cytoplasm of a tube-like cellular process (a neuron, a membrane nanotube or our parenchymal cells) includes (1) commonly considered viscous drag due to high cytoplasmic viscosity, (2) additional drag forces due to induced two-dimensional surface flow in the highly conformal membrane tube in close contact with the traveling cargo, and (3) an additional elastic energy term associated with net membrane deformation necessary to accommodate a large cargo. These multiple drag terms add up to produce additional dissipation, significantly slowing down a traveling rigid particle (such as a symbiont) inside the membrane tube.

Although the traveling particle inside the tube is almost two orders of magnitude larger than the tube dimensions, due to steric repulsion present because of entropic effects at finite non-zero temperatures (membrane fluctuations), a finite gap still exists between the symbiont and the associated outside membrane of the host cell. This gap was analytically estimated by Daniels [86] on average to be 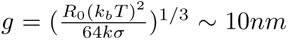 (for values previously discussed). In our FIB-SEM datasets (Fig.3), this gap corresponds to roughly 35nm and provides the key channel where cytoplasm can flow backwards around the symbiont, imparting it with an overall positive mobility.

In the above description, we do not include particle deformation (dinoflagellates are significantly stiffer than host cell due to its theca) since we do not observe any deformation of the symbiont during traversal (SI Videos 1,2,3). Assuming that the length of the host cell and associated membrane tension do not change significantly during symbiont movement, the host membrane close to the traveling symbiont must flow and bend to accommodate the large symbiont. Comparing the drag increase to traditional drag coefficient of 6*πηR*_0_ in a particle of size *R*_0_ embedded in Newtonian fluid of viscosity *η*, the increase in total drag has been predicted to be 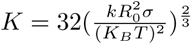, where membrane tension *σ* and bending rigidity *k* correspond to properties of the host cell [86]. Accounting for a 10*µm* symbiont in a membrane tube with typical membrane tension of 10^−4^*Jm*^−2^, we arrive at drag force ratio increase of a million fold higher than a cell freely moving in a cytoplasm of the same viscosity. The fact that symbionts achieve speeds higher than other reported actin-driven systems while overcoming such high drag forces is remarkable.

Extreme confinement might explain this paradox. Visually, dinoflagellates inside host cells resemble robbers from childhood cartoons, their head stretching out the stocking that covers them. The high degree of confinement of the thin host cell exacerbated by pressure exerted by surrounding large vacuolated cells essentially renders our system one-dimensional. Since actin filaments are 7nm in diameter and up to several micrometers in length[88], such high confinement can enable a high degree of alignment along the long axis of the cell, making it easier for actin to nucleate and generate a directional force on either side of the dinoflagellate, propelling them forward. This confinement of dense actin filaments is potentially similar in architecture to 1D actin-filled filopodia that exert forces on the membrane tips [89].

Further, in vitro assays have shown higher compressive forces led to increased filament density in Arp2/3-mediated growing branched actin networks[90]. Co-presence of myosin has also been suggested to synergize with Arp2/3 to enhance the efficiency of pushing forces in branched actin networks, reducing the density of actin network needed to generate propulsion forces[79]. Thus, confinement and higher load due to increased drag can increase force generation in actin-driven mobility in tubular structures such as the thin host cells in *Waminoa sp.*. This provides a plausible explanation for the paradoxical feat of dinoflagellates in our system traveling with high velocity against a high drag force. Taken together, we propose that symbionts in our system are driven by confinement-enhanced, Arp2/3 and myosin mediated, actin polymerization forces.

### 2.8 Symbiont distribution during diel rhythms and regeneration

Diel rhythm-based clustering of plastids is well-known intracellularly within dinoflagellates[91, 92]. Next we explore if, in a “Russian-doll-like” fashion, we see any diel rhythms in dinoflagellate distribution within the acoel worm’s tissue. We use macro-photography to image six worms cultured normally under 12h-12h light-dark cycle over a week (Fig.5A, SI Fig. 4). Curiously, we indeed observe a robust phenotype where symbionts are homogeneously distributed within host tissue during daytime, and at nighttime, symbionts retreat from the periphery of the worm and form smaller clusters throughout the host’s body (Fig.5A, SI Fig.4). This nighttime phenotype often leaves a peripheral ring devoid of symbionts in the worm (Fig.5A, SI Fig.4). To further probe whether the diel-rhythm is circadian, we cultured worms under 24 hour illumination and imaged them over a week (SI Fig.5). We still observe changes between homogeneous distribution and clustering of symbionts, but the day-night nature of this clustering is disrupted (SI Fig.5), most likely due to breakdown of circadian rhythms. This suggests that other signaling mechanisms are also present to create host-wide clustering events of symbionts.

**Fig. 5.**
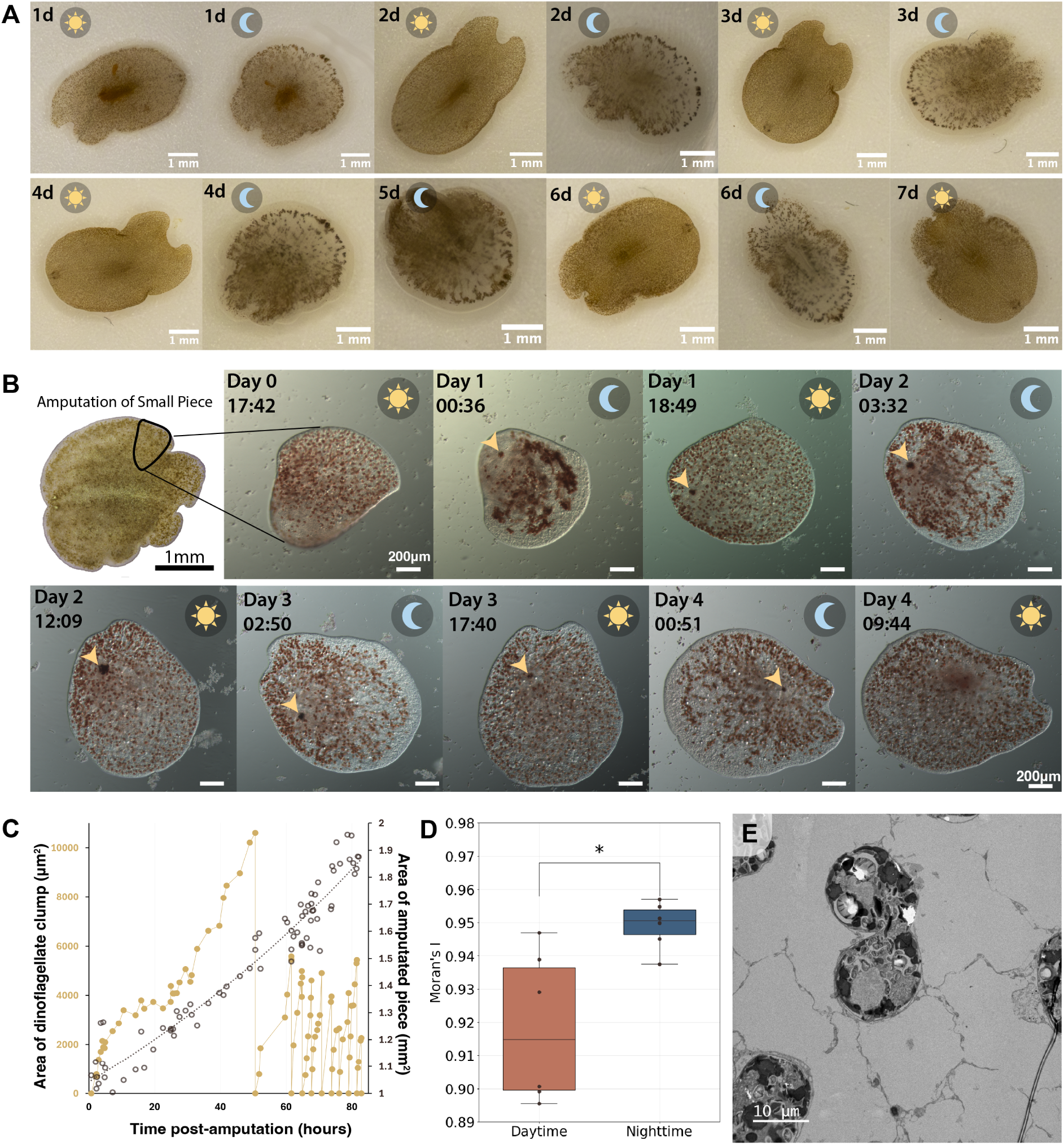
Long term imaging and regeneration experiments reveal diel rhythms in symbiont distribution and symbiont-clump formation during regeneration. (**A**) Using macro-photography of whole worms from natural populations cultured in standard conditions (12hour-12hour light-dark cycle, worms fed on Day 0 (0d), see Methods for details), we show that diel rhythms occur regularly in symbiont distribution (n=6, SI Fig.4). Specifically, we observe homogeneous symbiont distribution within host tissue during daytime, and at nighttime, symbionts retreat from the periphery of the worm and form smaller clusters throughout the host’s body. When cultured under 24 hour illumination, cyclic clumping is still observed, albeit no longer neatly aligned to day-night rhythms (n=6, SI Fig.5) (**B**) We amputate a small, initially mouthless piece of a worm, observing via tracking microscopy as it regenerates over the course of multiple days. We note the cyclic sequestration of symbionts into clumps (yellow arrowheads) followed by expulsion during the course of regeneration. Similar to in natural populations, cyclic clumping of symbionts at nighttime (moon icons) is also observed in the regenerating worm. (**C**) Quantifying both the size of the “symbiont clump” and the size of the regenerating worm, we note that the worm grows in size despite not having captured any external prey. The sequestration and expulsion of symbionts continues to occur even after the regeneration of the mouth. (**D**) Using Moran’s I as a metric, we find that symbionts are significantly more clustered at night as compared to during daytime for the regenerating worm in (B). ∗ : 0.01 *< p <* 0.05 (**E**) Dinoflagellates can also be observed undergoing cell division cycles within the worm.

Another remarkable attribute for which acoel worms have been studied is their capacity to regenerate whole worm morphology from amputated chunks[93]. We utilize this capacity to explore the role of symbiont mobility during regeneration, as it is a highly dynamic, animal-scale process. To explore this, we cut a small piece of tissue from an adult worm which does not include the mouth and thus cannot capture external prey (see Methods for details). We perform in-toto tracking microscopy of the amputated piece over the course of a week as it regenerates (SI Video 7, Fig.5B). Curiously, in the regenerating amputated piece, we observe a similar diel rhythm as we do in whole, normally cultured worms: more even distribution throughout the worm during daytime, and retreat from the worm’s perimeter combined with clustering at nighttime (Fig.5B,D). Interestingly, we also observe that a single clump of symbionts appears a few hours into the regeneration process (SI Video 7, yellow arrows in Fig.5B). When the worm regenerates a mouth after about two days, it expels the clump, and a new clump begins to form (SI Video 7, Fig.5C). In the expelled clump (SI Video 7), many dinoflagellates appear fragmented, suggesting cell lysis or death. This cycle of clump formation and expulsion continues well into the regeneration process, as the amputated piece’s area continues to increase without ingestion of any live prey (Fig.5C). We also find that symbionts go through division cycles inside host cells (Fig.5E) and thus increase in numbers over time. Furthermore, when worms are incubated in the dark, we see some evidence of symbionts being moved into the surrounding vacuolated cells (SI Fig.6). Most of these observations suggest the possibility that symbionts are being used as a food source via digestion by the host during the period when the organism does not have a mouth. Symbiont digestion by the host has also been previously reported as a possibility in corals [94], and “feeding spots” have been reported in another acoel worm, *C. longifissura*[93]. Taken together, we conjecture that symbiont digestion may be used for dinoflagellate population control, and the symbiont clumps observed during regeneration could be digested symbionts, though these hypotheses warrant further investigation.

## 3 Discussion

We discover organismal-scale, host actin-driven symbiont mobility in *Waminoa sp.*. This long-range mobility enables a highly dynamic spatiotemporal distribution of symbionts throughout the animal’s body, which manifests itself in diel rhythms and during regeneration. Our work challenges the longstanding notion of sessile symbionts in animal-microbe symbiosis and invites new ways of thinking about how holobionts maintain a dynamic homeostasis. Not only are symbionts mobile in *Waminoa sp.*, our findings suggest the host and/or dinoflagellates themselves may be able to sense relative positions and spatial distribution of the symbionts in the animal, enabling homogeneous or clustered distribution. This sensory ability could be manifested in the retained flagella [95], which is a fascinating direction for future investigation.

The retention of flagella could also indicate that our system represents a recent symbiosis [60]. Our study is a rare example of high-resolution in-vivo imaging of animal-microbe symbiosis over long timescales. It is possible that host-mediated spatiotemporal control of symbionts may be much more general and widespread as a phenomenon even in other systems where symbionts are currently considered to be sessile.

Our study also highlights the potentially widespread importance of host-mediated symbiont mobility in thinking about holobionts. Our finding of diel rhythms in symbiont distribution patterns finds interesting parallels in other organisms. For instance, glass frogs remove 89 percent of their red blood cells from circulation and pack them in their liver to appear transparent[96]. Scintillons and chloroplasts in the bioluminescent dinoflagellate *Pyrocystis lunula* migrate daily, with scintillons occupying the cell periphery at night and choloroplasts during the day [91, 92]. *P. lunula* can also contract its chloroplasts during the day for protection from strong light [97]. We’ve known for a long time that organisms maintain motile populations of cells such as macrophages. A similarly dynamic strategy is present in organelles being displaced throughout the cell. In single cells, examples of motile and mobile symbionts are both known, fulfilling important functions such as metabolic connection[28] and photoprotection[29] in their respective holobionts. Our discovery of host-mediated spatiotemporal control of symbiont distribution shows that in the animal world as well, symbiont mobility is much more than an earmark of individuality. Spatiotemporal control of symbionts within a tissue or a niche can benefit the holobiont via physically distributing metabolite exchange between the partners and may be a heretofore missing key piece in the regulation of symbiont populations and evolution of animal-microbe symbioses.

## 4 Methods

### 4.1 Worms and worm culture

Worms were kindly gifted by the curious aquarium hobbyists (Steven, Spensa, Cameron, and James) at Neptune Aquatics (San Jośe, CA). From these worms, we established two clonal lines that have been grown continuously. We verified using 18s and 28s sequencing that indeed we have an acoel worm species (*Waminoa sp.*) and they contain the well-known symbiont (*Amphidinium sp.*). Worms were cultured in 25mm deep petri dishes (VWR, 89107-632) in filtered natural seawater (FNSW) from Neptune Aquatics. Water parameters include (from Neptune Aquatics) - pH: 8.2-8.3, alkalinity: 10-11 (enhanced), calcium: 350-450 (enhanced), salinity: 1.024-1.026. Every 2 weeks, water is changed by transferring worms into new dishes filled with fresh FNSW using transfer pipette, then worms are fed freshly hatched brine shrimp. Petri dishes containing worms are kept on metro shelves that are lit by Fluval Marine 3.0 LED light panels, with a 12 hour-12 hour day-night cycle. We also observe that if some worms in the dish die, it may lead to all worms in the dish to die. To avoid complete culture crash, a backup culture is kept in a 20 gallon aquarium, set up with a water filter and the same lighting conditions as culture dishes. Worms in the backup tank are fed freshly hatched brine shrimp weekly.

### 4.2 Live tracking microscopy experiments

Worms were placed in glass bottom dishes (Cellvis, P06-1.5H-N), and allowed to settle on the microscope for 30 minutes up to a few hours without any disturbance. We notice that even a small vibration of the stage or the glass bottom dish can perturb a settled worm, and it starts moving again. When worms have settled, imaging was done is started under various live imaging modalities including a Nikon Eclipse Ti2 inverted microscope using Nikon 40x oil objective unless otherwise noted.

Unless otherwise noted, we use worms that have not been recently fed. This is because we find that transferring recently fed worms can result in tissue rupture on the dorsal side. Also unless otherwise noted, we do not use any muscle relaxants such as *MgCl*_2_ because we find that following *MgCl*_2_ treatment, worms tend to slightly detach from the surface, making microscopy videos more susceptible to effects of the worm jittering due to vibrations. It is also more difficult to image certain planes of interest in *MgCl*_2_-treated worms due to the detachment.

The longest single dinoflagellate track we observed (Fig.1F, SI Video 2) is an underestimate of the maximum possible distance a dinoflagellate can travel because the limit we report is due to a limit in our ability to track a dinoflagellate, rather than an observed boundary. Our tracks of single dinoflagellates end either when dinoflagellates move out of focus (since the “highway system” is 3-dimensional in nature) or when the worm moves, and we lose track of the dinoflagellate.

In tracks obtained, animals are not always completely still. For comparing dinoflagellate and vesicle mobility across different planes of the worm, only cases where stable tracks were successfully obtained for both ventral and mid networks at approximately the same spot on the worm were included for further analysis to rule out potential data skew due to different regions of worm. Segments where the animal is still are selected, and then Cellpose-Trackmate[98, 99] was used to obtain tracks of dinoflagellates and vesicles.

The data in Fig.2 represents cumulative results from a total of 8 different spots imaged over worms. The total n was 144 dinoflagellates in the mid layer, 658 vesicles in the mid layer, 117 dinoflagellates in the ventral layer, and 994 vesicles in the ventral layer. Detailed list of number of trials and segments, as well as number dinoflagellates and vesicles from each trial, can be found at https://www.notion.so/Nighthawk-tracking-number-of-trials-112fc95b01b180968d97cf6297d49a7b?pvs=4. Mann-Whitney U Test was used to determine significance. For quantifying velocity, we choose to use mean momentary velocity (sliding window analysis with overlapping windows of 6 seconds) as a metric because it is more resilient than instantaneous velocity to potential effects of jitter. For calculation of slope of log-log MSD vs delta t, only tracks with no gaps are used. MSD is calculated using delta t up to 90 seconds, and a linear best fit was done on the log-log plot of MSD vs delta t.

### 4.3 Live confocal imaging of actin dynamics

For live confocal imaging, similar to DIC microscopy experiments, we use worms that have not been recently fed. However, there are a few key differences in protocol, so we did not include mobility data from DIC microscopy experiments in the “no drug” condition (i.e. no overlap of data reported in Fig.2G-H and Fig.4G-H), to avoid any potential data skew of mobility differences that may be caused by these differences in protocol.

For live confocal imaging of actin dynamics, the results of which are presented in Fig.4, we first pipette 5 worms into a Tomodish (901002-02), which is used due to availability, but can be replaced by any glass-bottom dish that has a shallow well and fits well onto the confocal setup. We first prepare in a 1mL Eppendorf tube a solution of 2.5*µL* of CellMask Orange Actin Tracking Stain (Invitrogen A57247) added to 995uL of fresh FNSW. For experiments with Hoechst and DiD, we also added 1:10000 Hoechst 33342 and 2.5*µL* DiD stain (Thermofisher scientific, V22887). We then pipette out the seawater in the TomoDish (worms will survive this because they secrete a coating that protects them against short periods of exposure to air), and replace it with the freshly prepared solution. Then, we use a single-use surgical scalpel to slice the worms into a “head” piece and a “tail piece”. We pipette out the head pieces and keep the tail pieces in the dish. The reason we choose to put 5 indistinguishable worms on one dish is so that we are more likely to have some worms that are sitting still at any given point in time. We slice them in half and keep the tail piece for two reasons. Firstly, we note that worms are very sensitive to and become highly motile under laser illumination, making confocal imaging very difficult. Thus, we find that when sliced in half, the tail piece is much less active and enables long term live confocal imaging to be performed on the worm. Secondly, we cut the worms for easy dye penetration which allows us to observe actin motility live, which is crucial for this experiment. To try to avoid potential data skew from injury, we image near the tail (further from injury site). We use an inverted Zeiss LSM780 laser scanning confocal microscope (Stanford, CSIF imaging facility).

Because in each experiment, the five worms are indistinguishable, we repeat each condition over at least five different experiments, tracking dinoflagellates from up to five worms per experiment. For investigating the effect of Latrunculin A on live actin dynamics, we added 10*µL* of 1mM Latrunculin A (Sigma-Aldrich, 428026) to the prepared solution (2.5*µL* of CellMask Orange Actin Tracking Stain (Invitrogen A57247) in 995*µL* of fresh FNSW). For control experiments, we added 10*µL* of DMSO instead.

Similar to the data processing pipeline for DIC experiments, segments where the animal is still are selected, and then Cellpose-Trackmate[98, 99] was used to obtain tracks of dinoflagellates and vesicles. This was done using autofluoresence images collected using Alexa Fluor 647 filter settings for dinoflagellates, and transmitted images for vesicles. Because animals are jittery under laser illumination and it is difficult to obtain still tracks, some sequences had minor drift and were first stabilized using the Image Stabilizer plugin in Fiji[100]. Transmitted images from confocal microscopy is more difficult to use than DIC microscopy for tracking vesicles, so we have less successful tracks of vesicles. The total number of dinoflagellates (over a total of five independent experiments) represented for each condition presented in Fig.4G-H are: 972 dinoflagellates from 7 experiments tracked in ventral layer (no drug), 1015 dinoflagellates from 5 experiments tracked in mid layer (no drug), 629 dinoflag-ellates from 5 experiments tracked in mid layer (LatA), 512 dinoflagellates from 5 experiments tracked in ventral layer (LatA), 930 dinoflagellates from 5 experiments tracked in ventral layer (DMSO), and 575 dinoflagellates from 5 experiments tracked in mid layer (DMSO). The total numbers of vesicles tracked are 121 vesicles from 2 experiments tracked in ventral layer (no drug), 56 vesicles from 2 experiments tracked in mid layer (no drug), 82 vesicles from 3 experiments tracked in mid layer (LatA), 179 vesicles from 2 experiments tracked in ventral layer (LatA), 10 vesicles from 1 experiment tracked in mid layer (DMSO), 12 vesicles from 1 experiment tracked in ventral layer (DMSO). Detailed list of number of trials and segments, as well as number dinoflagellates and vesicles from each trial, can be found at https://www.notion.so/Confocal-Tracking-number-of-trials-and-days-10efc95b01b1805d9b65d95f7ddfd424?pvs=4. Mann-Whitney U Test was used to determine significance.

### 4.4 Temporal dynamics of drug inhibition

We investigated the temporal dynamics of the mobility phenotype following ML-7, LatA, CK-666, Blebbistatin, and Nocodazole treatment. For ML-7, we dissolved 1mg of ML-7 (Sigma Aldrich, 475880) in DMSO to make a 10mM stock solution. Worms were placed in glass bottom dishes (Cellvis, P06-1.5H-N), filled with 10mL of FNSW each, and allowed to settle overnight. DIC imaging of worms prior to drug treatment was done on Nikon Eclipse Ti2 inverted microscope using Nikon 40x oil objective unless otherwise noted. 2*µL* of 10mM ML-7 solution was added for a final concentration of 2*µM*, and DIC iamging was again done under the same settings. For the effect of low-concentration LatA, we used the same protocol, instead adding 2*µL* 1mM LatA (for final LatA concentration of 200nM). For Nocodazole experiments, we used the same protocol, instead adding 1.5*µL* of 20mM Nocodazole, for a final concentration of 3*µM*. For CK-666 experiments, we dissolved 25mg of CK-666 in 843uL DMSO to make a 100mM stock and tried final concentrations of 50*µM*, 30*µM*, and 10*µM*, shown in SI Video 6. CK-666 causes worms to be overly active and difficult to image, so we were unable to quantify the data and show video from 1 worm for each of 3 concentrations in SI Video 6. For Blebbistatin, we added 2*µL* of a 50mM stock ready-made stock in DMSO (Sigma-Aldrich 203389) for a final concentration of 10uM.

We were able to quantify results for ML-7, LatA, Blebbistatin, and Nocodazole. For Nocodazole (Fig.4A), LatA (Fig.4B), and Blebbistatin (SI Fig. 2), we have data for 3 worms each. For ML-7 (Fig.4C), we have data for 6 worms. Because we cannot control when the worms would be still, we did not have fixed time points across the trials. Instead, we imaged at various time points for different worms and collated the data to result in Fig. 4A-C, where we use different markers to show results from different worms. We show the top 50 percent fastest moving dinoflagellates at each time point. Details on number of repeats for each drug experiment is found at https://www.notion.so/Nighthawk-latA-ML7-Nocodazole-number-of-trials-858d266275164d7fb6424136c42d49d8?pvs=4.

Similar to the data processing pipeline for other DIC experiments, segments where the animal is still are selected. For plots in Fig.4, segments in the mid network were selected for further data processing. Because animals are jittery following drug addition, often even the most still segments we can select have minor drift. Thus, segments are first stabilized using the Image Stabilizer plugin in Fiji[100]. Then Cellpose-Trackmate[98, 99] was used to obtain tracks of dinoflagellates.

### 4.5 Electron microscopy

For TEM images presented in Fig. 5 and Supplementary Information, worms were fixed in 2.5% gluteraldehyde and 2% paraformaldehyde, added to concentrated artificial seawater to result in a final salinity 35ppt, or specific gravity of 1.026. Samples were then incubated in 1% Osmium tetroxide + 1.6% Potassium ferricyanide in seawater for 1 hour on a rocker, followed by acetone dehydration and resin infiltration, outlined in more detail at https://em-lab.berkeley.edu/EML/tem-generic-protocol. We then performed TEM imaging on a FEI Tecnai 12 transmission electron microscope. We note that fixation in seawater is crucial for preserving cellular features and avoiding artifacts.

For TEM images and FIB-SEM data presented in Fig.3, as well as SEM images presented in Fig.2, we used worms that were fixed using the high pressure freezing protocol outlined in Gallet et al. 2004 [101]. Briefly, high pressure freezing machine (Leica system) is used to cryofix the worms, followed by freeze substitution and resin embedding. The sample is then processed for SEM, TEM, and FIB-SEM respectively.

Focused Ion beam Scanning Electron Microscopy (FIB-SEM) tomography was performed with a Zeiss CrossBeam 550 microscope (Zeiss, Germany) as in Uwizeye et al 2021 and Rao et al 2025 https://www.cell.com/current-biology/fulltext/S0960-9822(25)00392-6. The resin block containing the cells was fixed on a stub with silver paste and surface abraded with a diamond knife in a microtome to obtain a flat and clean surface. Samples were then metallized with 4–8 nm of platinum to avoid charging during observations. Inside the FIB-SEM, a second platinum layer (1–2 mm) was deposited locally on the analyzed area to mitigate possible curtaining artifacts. The sample was then abraded slice by slice with the Ga+ ion beam (typically with a current of 700 pA at 30 kV). Each exposed surface was imaged by SEM at 1.5 kV and with a current of 1 nA using the in-column EsB backscatter detector. Similar milling and imaging modes were used for all samples. Automatic focus and astigmatism correction were performed during image acquisition, typically at approximately hourly intervals. For each slice, a thickness of 10nm was removed and SEM images were recorded with a corresponding pixel size of 10nm in order to obtain an isotropic voxel size. Entire volumes were imaged with 800–3000 frames for the worm. The first steps of image processing were performed using Fiji software for registration (adapted StackReg plugin), and noise reduction (3D mean function of the 3D suite plugin).

The voxel size for FIB-SEM is 10nm by 10nm by 10nm, but due to file size, for further processing, subsampling was done so that voxel size is 20nm by 20nm by 20nm. Segmentation is done manually layer by layer on Dragonfly software, so the noodle-like texture in the host cells is an artifact. Segmentation is done by manually tracing features known to a specific cell type or organelle across the layers (for instance, characteristic morphologies of dinoflagellates and nuclei in EM, striation for muscle cells, cells hosting dinoflagellates by tracing the cytoplasm that surrounds the dinoflagellate).

### 4.6 Long term tracking microscopy of regeneration process

For long term tracking experiments during regeneration, we amputated a small piece from the worm near the tail region. Animals were placed in a well within a glass-bottom 6-well plate (Cellvis P06-1.5H-N) and observed under white LED illumination on a Squid tracking microscope[50] (Cephla) with 10x magnification and Imaging Source DFK 33UX226 4000x3000 pixel camera. The well was filled to the top (around 11mL) with filtered seawater and covered with a large glass slide to prevent evaporation. One single newly hatched brine shrimp was added to the dish prior to starting the experiment. The worm did not eat the shrimp, confirming that the piece we cut does not contain the animal’s mouth. Water changes were done on Day 2, Day 3, and Day 5, wherein 3 transfer pipettes (around 4.5mL) of water was removed from the dish and replaced with fresh filtered seawater. A small number (around 3) freshly added brine shrimp were added at each water change. For calculation of area of dinoflagellate clump and area of worm, clumps and worms were manually outlined on Fiji and the areas determined. Moran’s I was calculated for 6 images during day time and 6 images during night time (1 image per day post amputation). Mann-Whitney U Test was used to determine significance.

### 4.7 Low magnification imaging of uncut worms

For day-night imaging of whole, uncut worms to discern if a) the diel rhythm in symbiont distribution observed in the regenerating worm also occurs naturally, and b) whether the diel rhythm observed is circadian or light-condition-dependent, we use macro-photography, since worms are too large to wholly fit under magnified microscopy. 12 worms were placed in 2 6-well glass bottom dishes (Cellvis, P06-1.5H-N). On Day 0, one dish was moved out of the incubator to be cultured under perpetual white light, the other continued to be cultured normally in the incubator under 12 hour light-dark cycle (illuminated 7:00-19:00). On Day 0, animals were also fed brine shrimp. On Day 1, water change was performed to remove remaining brine shrimp and the photo series was started. At various times of day, both dishes were imaged in a foldable photo box (Amazon) using a Fujifilm XT-4 mirrorless camera with a XF80mmF2.8 R LM OIS WR Macro lens. A 0.25 inch green sticker (parts of which show up in some images presented) is used in each photo for scale. RAW files were captured and denoised in Adobe Lightroom, then scale bar is added (using the sticker as a reference), and image is cropped in Fiji.

## Supporting information

SI Video 1 - Symbiont mobility DIC movie

SI Video 2 - Three party symbiosis

SI Video 3 - Symbiont mobility in morphologically distinct layers of the worm

SI Video 4 - FIB-SEM-based 3D reconstruction

SI Video 5 - Live actin staining

SI Video 6 - Drug assays

SI Video 7 - Symbiont clump formation in regeneration

## Supplementary information

- Supplementary Information PDF - Contains supplementary figures
- SI Movie 1 - DIC microscopy movie showing symbiont mobility in the tissue of the *Waminoa sp.* acoel worm
- SI Movie 2 - Introduction to the “three party symbiosis”
- SI Movie 3 - DIC microscopy movies showing symbiont mobility in two morphologically distinct layers (mid and ventral) of the worm
- SI Movie 4- Segmented FIB-SEM data showing symbionts’ retention of flagella, intracellular nature of symbionts, and host tissue structure surrounding symbionts
- SI Movie 5 - Live actin staining shows flowing actin puncta in cells hosting dinoflagellates
- SI Movie 6 - Drug assays (LatA, CK-666, ML-7) confirm primary role of host actin in symbiont mobility
- SI Movie 7 - Long term imaging of a regenerating worm reveals symbiont clump formation during regeneration

## Acknowledgments.

We thank the curious folks (especially Steven, Spensa, Cameron, and James) at Neptune Aquatics for gifting us with worms and for worm care advice. We thank the staff at the University of California Berkeley Electron Microscope Laboratory - especially Danielle Jorgens and Reena Zalpuri - for advice and assistance in electron microscopy sample preparation and data collection. We thank the staff at the Cell Sciences Imaging Facility (CSIF) at Stanford - especially Ruth Yamawaki, John Perrino, and David Lenzi for their help and advice with EM and confocal microscopy. We thank Rebecca Konte for illustration of the schematic and artistic input on the figures. We thank Dania Nanes-Sarfati, Robin Bigasin, and Nelson Hall for advice on worm care. We thank Qing Zhang, Andrew Guan, Nelson Hall, and Magnus Bauer for help with worm maintenance when we were away from lab. G.Z. is grateful to committee members and all Prakash Lab members for suggestions and stimulating discussions. Part of this work was performed at the Stanford Nano Shared Facilities (SNSF), supported by the National Science Foundation under award ECCS-2026822. Part of this work was performed at the CSIF at Stanford. Research was supported by the AtlaSymbio project funded by the Gordon and Betty Moore Foundation (GBMF11532, https://doi.org/10.37807/GBMF11532). FIB-SEM and SEM imaging were carried out on the Platform for Nanocharacterization (PFNC) of IRIG CEA-Grenoble, supported by the “Recherche Technologique de Base” and “France 2030—ANR-22-PEEL-0014” programs of the French National Research Agency (ANR). We thank Benoit Gallet from the IBS/ISBG EM platform, part of the Grenoble Instruct-ERIC center (ISBG; UMS 3518 CNRS-CEA-UGAEMBL), within the Grenoble Partnership for Structural Biology (PSB), which is supported by FRISBI (ANR-10-INBS-05-02) and GRAL, financed within the University Grenoble Alpes graduate school (Ecoles Universitaires de Recherche) CBH-EUR-GS (ANR-17-EURE-0003). The IBS/ISBG EM facility is led by Dr. Guy Schoehn and supported by the Auvergne Rhone-Alpes Region, the Fondation Recherche Medicale (FRM), the fonds FEDER, and the GIS-Infrastructures en Biologie Sante et Agronomie (IBISA). We acknowledge support from Schmidt Futures Innovation Fellowship (M.P.), Moore Foundation Research Grant (M.P.), NSF CCC (DBI1548297 (M.P.)) grant, Woods Institute for the Environment (M.P), Stanford DARE doctoral fellowship (G.Z.), and NSERC PGS-D (G.Z.).

## Author contributions

G.Z. and M.P. conceptualized the study. G.Z. conducted the experiments except SEM, TEM, and FIB-SEM performed using high-pressure-frozen worms. G.Z. analyzed the data in discussion with M.P. M.P. contributed theoretical calculations. G.T., P-H J., and J.D. performed SEM, TEM, and FIB-SEM using high-pressure-frozen worms. G.Z. and M.P. drafted the manuscript. All authors edited the manuscript.

## References

[1] Margulis, L. Symbiotic Planet: A New Look At Evolution (Basic Books, London, England, 1999).

[2] Gilbert, S. F., Sapp, J. & Tauber, A. I. A symbiotic view of life: we have never been individuals. Q. Rev. Biol. 87, 325–341 (2012).

[3] Barott, K. L., Venn, A. A., Perez, S. O., Tambutté, S. & Tresguerres, M. Coral host cells acidify symbiotic algal microenvironment to promote photosynthesis. Proc. Natl. Acad. Sci. U. S. A. 112, 607–612 (2015).

[4] Gruber-Vodicka, H. R. et al. Paracatenula, an ancient symbiosis between thiotrophic alphaproteobacteria and catenulid flatworms. Proc. Natl. Acad. Sci. U. S. A. 108, 12078–12083 (2011).

[5] Douglas, A. E. Housing microbial symbionts: evolutionary origins and diversification of symbiotic organs in animals. Philos. Trans. R. Soc. Lond. B Biol. Sci. 375, 20190603 (2020).

[6] Kaur, R. et al. Living in the endosymbiotic world of wolbachia: A centennial review. Cell Host Microbe 29, 879–893 (2021).

[7] Gruber-Vodicka, H. R. et al. Two intracellular and cell type-specific bacterial symbionts in the placozoan trichoplax H2. Nat. Microbiol. 4, 1465–1474 (2019).

[8] Militello, G. Motility control of symbionts and organelles by the eukaryotic cell: The handling of the motile capacity of individual parts forges a collective biological identity. Front. Psychol. 10, 2080 (2019).

[9] Davy Simon K., Allemand Denis & Weis Virginia M. Cell biology of cnidarian-dinoflagellate symbiosis. Microbiol. Mol. Biol. Rev. 76, 229–261 (2012).

[10] Jimbo, M. et al. Possible involvement of glycolipids in lectin-mediated cellular transformation of symbiotic microalgae in corals. J. Exp. Mar. Bio. Ecol. 439, 129–135 (2013).

[11] Nyholm, S. V. & McFall-Ngai, M. J. The winnowing: establishing the squid-vibrio symbiosis. Nat. Rev. Microbiol. 2, 632–642 (2004).

[12] Nyholm, S. V. & McFall-Ngai, M. J. A lasting symbiosis: how the hawaiian bobtail squid finds and keeps its bioluminescent bacterial partner. Nat. Rev. Microbiol. 19, 666–679 (2021).

[13] Cullender, T. C. et al. Innate and adaptive immunity interact to quench microbiome flagellar motility in the gut. Cell Host Microbe 14, 571–581 (2013).

[14] Akahoshi, D. T. & Bevins, C. L. Flagella at the host-microbe interface: Key functions intersect with redundant responses. Front. Immunol. 13, 828758 (2022).

[15] Yee, D. P. et al. Physiology and metabolism of eukaryotic microalgae involved in aquatic photosymbioses. New Phytol. (2025).

[16] Ruiz-Trillo, I., Riutort, M., Littlewood, D. T., Herniou, E. A. & Bagunã, J. Acoel flatworms: earliest extant bilaterian metazoans, not members of platyhelminthes. Science 283, 1919–1923 (1999).

[17] Philippe, H. et al. Acoelomorph flatworms are deuterostomes related to xenoturbella. Nature 470, 255–258 (2011).

[18] Álvarez Presas, M., Ruiz-Trillo, I. & Paps, J. Novel genomic approaches support xenacoelomorpha as sister to all bilateria. Research Square (2024).

[19] Keeble, F. Plant-animals; a study in symbiosis (Cambridge University Press, 1910).

[20] Junker, A. D., Chen, J. Z., DuBose, J. G. & Gerardo, N. M. Dynamic reciprocal morphological changes in insect hosts and bacterial symbionts. J. Exp. Biol. 228, jeb249474 (2025).

[21] Millikan, D. S. & Ruby, E. G. Vibrio fischeri flagellin a is essential for normal motility and for symbiotic competence during initial squid light organ colonization. J. Bacteriol. 186, 4315–4325 (2004).

[22] Koike, K. et al. Octocoral chemical signaling selects and controls dinoflagellate symbionts. Biol. Bull. 207, 80–86 (2004).

[23] Bhattacharya, D., Stephens, T. G., Chille, E. E., Benites, L. F. & Chan, C. X. Facultative lifestyle drives diversity of coral algal symbionts. Trends Ecol. Evol. 39, 239–247 (2024).

[24] LaJeunesse, T. C. Zooxanthellae. Curr. Biol. 30, R1110–R1113 (2020).

[25] Ruby, E. G. & Asato, L. M. Growth and flagellation of vibrio fischeri during initiation of the sepiolid squid light organ symbiosis. Arch. Microbiol. 159, 160–167 (1993).

[26] Billman, G. E. Homeostasis: The underappreciated and far too often ignored central organizing principle of physiology. Front. Physiol. 11, 200 (2020).

[27] Baker, A. C., Starger, C. J., McClanahan, T. R. & Glynn, P. W. Coral reefs: corals’ adaptive response to climate change. Nature 430, 741 (2004).

[28] Decelle, J. et al. Subcellular architecture and metabolic connection in the planktonic photosym-biosis between collodaria (radiolarians) and their microalgae. Environ. Microbiol. 23, 6569–6586 (2021).

[29] Petrou, K., Ralph, P. J. & Nielsen, D. A. A novel mechanism for host-mediated photoprotection in endosymbiotic foraminifera. ISME J. 11, 453–462 (2017).

[30] Feng, Q. & Kornmann, B. Mechanical forces on cellular organelles. J. Cell Sci. 131, jcs218479 (2018).

[31] Venkatesh, K., Mathew, A. & Koushika, S. P. Role of actin in organelle trafficking in neurons. Cytoskeleton (Hoboken) 77, 97–109 (2020).

[32] Morris, R. L. & Hollenbeck, P. J. Axonal transport of mitochondria along microtubules and F-actin in living vertebrate neurons. J. Cell Biol. 131, 1315–1326 (1995).

[33] Ricci, L. & Srivastava, M. Transgenesis in the acoel worm hofstenia miamia. Dev. Cell 56, 3160–3170.e4 (2021).

[34] Duruz, J. et al. Acoel single-cell transcriptomics: Cell type analysis of a deep branching bilaterian. Mol. Biol. Evol. 38, 1888–1904 (2021).

[35] Reinhard M. Reiger, Seth Tyler, Julian P.S. Smith III, and Gunde E. Rieger. in Platyhelminthes: Turbellaria (ed. Frederick W. Harrison and Burton J. Bogitsh) Microscopic anatomy of invertebrates, Volume 3 7–141 (Wiley-Liss, New York, NY, 1991).

[36] Todt, C. Structure and evolution of the pharynx simplex in acoel flatworms (acoela). J. Morphol. 270, 271–290 (2009).

[37] Pedersen, K. J. The cellular organization of convoluta convoluta, an acoel turbellarian: A cytological, histochemical and fine structural study. Z. Zellforsch. Mikrosk. Anat. 64, 655–687 (1964).

[38] Creée, M. Monograph of the solenofilomorphidae (turbellaria: Acoela). Int. Rev. Gesamt. Hydrobiol. 60, 769–845 (1975).

[39] Zabotin, Y. I. & Golubev, A. I. Ultrastructure of oocytes and female copulatory organs of acoela. Biology Bulletin 41, 722–735 (2014).

[40] Tekle, Y. I., Raikova, O. I., Ahmadzadeh, A. & Jondelius, U. Revision of the childiidae (acoela), a total evidence approach in reconstructing the phylogeny of acoels with reversed muscle layers. J. Zoolog. Syst. Evol. Res. 43, 72–90 (2005).

[41] Hooge, M. D. Evolution of body-wall musculature in the platyhelminthes (acoelomorpha, catenulida, rhabditophora). J. Morphol. 249, 171–194 (2001).

[42] O. Barneah, I. Brickner, M. Hooge, V. M. Weis, T. C. LaJeunesse & Y. Benayahu. Three party symbiosis: acoelomorph worms, corals and unicellular algal symbionts in eilat (red sea). Mar. Biol. 151, 1215–1223 (2007).

[43] Maggioni, G. et al. The association of waminoa with reef corals in singapore and its impact on putative immune- and stress-response genes. Diversity (Basel) 14, 300 (2022).

[44] Cooper, C., Clode, P. L., Thomson, D. P. & Stat, M. A flatworm from the genus waminoa (acoela: Convolutidae) associated with bleached corals in western australia. Zoolog. Sci. 32, 465–473 (2015).

[45] Huang, C.-Y., Hwang, J.-S., Yamashiro, H. & Tang, S.-L. Spatial and cross-seasonal patterns of coral diseases in reefs of taiwan: high prevalence and regional variation. Dis. Aquat. Organ. 146, 145–156 (2021).

[46] Ogunlana, M. V. et al. Waminoa brickneri n. sp. (acoela: Acoelomorpha) associated with corals in the red sea. Zootaxa 1008, 1 (2005).

[47] Kunihiro, S. & Reimer, J. D. Phylogenetic analyses of symbiodinium isolated from waminoa and their anthozoan hosts in the ryukyu archipelago, southern japan. Symbiosis 76, 253–264 (2018).

[48] Haapkylä, J., et al. Association of waminoa sp. (acoela) with corals in the wakatobi marine park, south-east sulawesi, indonesia. Mar. Biol. 156, 1021–1027 (2009).

[49] Lopes, R. M. & Silveira, M. Symbiosis between a pelagic flatworm and a dinoflagellate from a tropical area: structural observations. Hydrobiologia 287, 277–284 (1994).

[50] Li, H., et al. Squid: Simplifying quantitative imaging platform development and deployment. bioRxiv 2020.12.28.424613 (2020).

[51] Achatz, J. G., Chiodin, M., Salvenmoser, W., Tyler, S. & Martinez, P. The acoela: on their kind and kinships, especially with nemertodermatids and xenoturbellids (bilateria incertae sedis). Org. Divers. Evol. 13, 267–286 (2013).

[52] Hanson, E. D. Regeneration in acoelous flatworms: The role of the peripheral parenchyma. Wilhelm Roux Arch. Entwickl. Mech. Org. 159, 298–313 (1967).

[53] Gavián, B., Sprecher, S. G., Hartenstein, V. & Martinez, P. The digestive system of xenacoelomorphs. Cell Tissue Res. 377, 369–382 (2019).

[54] Hikosaka-Katayama, T., Koike, K., Yamashita, H., Hikosaka, A. & Koike, K. Mechanisms of maternal inheritance of dinoflagellate symbionts in the acoelomorph worm waminoa litus. Zoolog. Sci. 29, 559–567 (2012).

[55] Chang, R. & Prakash, M. Topological damping in an ultrafast giant cell. Proc. Natl. Acad. Sci. U. S. A. 120, e2303940120 (2023).

[56] Smith, J. P. S., III. Fine-structural observations on the central parenchyma in convoluta sp. Hydrobiologia 84, 259–265 (1981).

[57] Oschman, J. L. Development of the symbiosis of convoluta roscoffensis graff and platymonas sp. J. Phycol. 2, 105–111 (1966).

[58] Hirose, E. & Hirose, M. Body colors and algal distribution in the acoel flatworm convolutriloba longifissura: histology and ultrastructure. Zoolog. Sci. 24, 1241–1246 (2007).

[59] Bailly, X. et al. The chimerical and multifaceted marine acoel symsagittifera roscoffensis: from photosymbiosis to brain regeneration. Front. Microbiol. 5, 498 (2014).

[60] Taylor, D. On the symbiosis between amphidinium klebsii [dinophyceae] and amphiscolops langerhansi [turbellaria: Acoela]. J. Mar. Biol. Assoc. U. K. 51, 301–313 (1971).

[61] Fenchel, T. How dinoflagellates swim. Protist 152, 329–338 (2001).

[62] Kumar, M. S. & Philominathan, P. The physics of flagellar motion of E. coli during chemotaxis. Biophys. Rev. 2, 13–20 (2010).

[63] Meddens, M. B. M. et al. Actomyosin-dependent dynamic spatial patterns of cytoskeletal components drive mesoscale podosome organization. Nat. Commun. 7, 13127 (2016).

[64] Coé, M., Brenner, S. L., Spector, I. & Korn, E. D. Inhibition of actin polymerization by latrunculin a. FEBS Lett. 213, 316–318 (1987).

[65] Saitoh, M., Ishikawa, T., Matsushima, S., Naka, M. & Hidaka, H. Selective inhibition of catalytic activity of smooth muscle myosin light chain kinase. J. Biol. Chem. 262, 7796–7801 (1987).

[66] Isemura, M., Mita, T., Satoh, K., Narumi, K. & Motomiya, M. Myosin light chain kinase inhibitors ML-7 and ML-9 inhibit mouse lung carcinoma cell attachment to the fibronectin substratum. Cell Biol. Int. Rep. 15, 965–972 (1991).

[67] Melkov, A. & Abdu, U. Regulation of long-distance transport of mitochondria along microtubules. Cell. Mol. Life Sci. 75, 163–176 (2018).

[68] Frederick, R. L. & Shaw, J. M. Moving mitochondria: establishing distribution of an essential organelle. Traffic 8, 1668–1675 (2007).

[69] Goldstein, L. S. & Yang, Z. Microtubule-based transport systems in neurons: the roles of kinesins and dyneins. Annu. Rev. Neurosci. 23, 39–71 (2000).

[70] Umeshima, H. et al. Local traction force in the proximal leading process triggers nuclear translocation during neuronal migration. Neurosci. Res. 142, 38–48 (2019).

[71] Umeshima, H., Hirano, T. & Kengaku, M. Microtubule-based nuclear movement occurs independently of centrosome positioning in migrating neurons. Proceedings of the National Academy of Sciences 104, 16182–16187 (2007).

[72] Theriot, J. A., Mitchison, T. J., Tilney, L. G. & Portnoy, D. A. The rate of actin-based motility of intracellular listeria monocytogenes equals the rate of actin polymerization. Nature 357, 257–260 (1992).

[73] Mogilner, A. & Oster, G. Cell motility driven by actin polymerization. Biophys. J. 71, 3030–3045 (1996).

[74] Jasnin, M. & Crevenna, A. H. Quantitative analysis of filament branch orientation in listeria actin comet tails. Biophys. J. 110, 817–826 (2016).

[75] Lacayo, C. I. & Theriot, J. A. Listeria monocytogenes actin-based motility varies depending on subcellular location: a kinematic probe for cytoarchitecture. Mol. Biol. Cell 15, 2164–2175 (2004).

[76] Murrell, M., Oakes, P. W., Lenz, M. & Gardel, M. L. Forcing cells into shape: the mechanics of actomyosin contractility. Nat. Rev. Mol. Cell Biol. 16, 486–498 (2015).

[77] Thiam, H.-R. et al. Perinuclear Arp2/3-driven actin polymerization enables nuclear deformation to facilitate cell migration through complex environments. Nat. Commun. 7, 10997 (2016).

[78] Shu, S., Liu, X. & Korn, E. D. Blebbistatin and blebbistatin-inactivated myosin II inhibit myosin II-independent processes in dictyostelium. Proc. Natl. Acad. Sci. U. S. A. 102, 1472–1477 (2005).

[79] Xu, M. et al. Myosin-I synergizes with Arp2/3 complex to enhance the pushing forces of branched actin networks. Sci. Adv. 10, eado5788 (2024).

[80] Rustom, A., Saffrich, R., Markovic, I., Walther, P. & Gerdes, H.-H. Nanotubular highways for intercellular organelle transport. Science 303, 1007–1010 (2004).

[81] Saha, T. et al. Intercellular nanotubes mediate mitochondrial trafficking between cancer and immune cells. Nat. Nanotechnol. 17, 98–106 (2022).

[82] He, K. et al. Intercellular transportation of quantum dots mediated by membrane nanotubes. ACS Nano 4, 3015–3022 (2010).

[83] Kolba, M. D. et al. Tunneling nanotube-mediated intercellular vesicle and protein transfer in the stroma-provided imatinib resistance in chronic myeloid leukemia cells. Cell Death Dis. 10, 817 (2019).

[84] Yang, X. et al. Mitochondrial dynamics quantitatively revealed by STED nanoscopy with an enhanced squaraine variant probe. Nat. Commun. 11, 3699 (2020).

[85] Dagar, S. & Subramaniam, S. Tunneling nanotube: An enticing cell-cell communication in the nervous system. Biology (Basel) 12, 1288 (2023).

[86] Daniels, D. R. Transport of solid bodies along tubular membrane tethers. PLoS One 14, e0210259 (2019).

[87] Kozlov, M. M., Weissenhorn, W. & Bassereau, P. in Membrane remodeling: theoretical principles, structures of protein scaffolds and forces involved 287–349 (Oxford University Press, 2016).

[88] Cooper, G. M. in Structure and organization of actin filaments (Sinauer Associates, 2000).

[89] Bornschlögl, T. How filopodia pull: what we know about the mechanics and dynamics of filopodia: Bornschlögl. Cytoskeleton (Hoboken) 70, 590–603 (2013).

[90] Li, T.-D., Bieling, P., Weichsel, J., Mullins, R. D. & Fletcher, D. A. The molecular mechanism of load adaptation by branched actin networks. Elife 11 (2022).

[91] Seo, K. S. & Fritz, L. Cell ultrastructural changes correlate with circadian rhythms in *Pyrocystis lunula* (pyrrophyta). J. Phycol. 36, 351–358 (2000).

[92] Heimann, K., Klerks, P. L. & Hasenstein, K. H. Involvement of actin and microtubules in regulation of bioluminescence and translocation of chloroplasts in the dinoflagellate *Pyrocystis lunula*. Botanica Marina 52, 170–177 (2009).

[93] Nanes Sarfati, D., et al. Coordinated wound responses in a regenerative animal-algal holobiont. Nat. Commun. 15, 4032 (2024).

[94] Wiedenmann, J. et al. Reef-building corals farm and feed on their photosynthetic symbionts. Nature 620, 1018–1024 (2023).

[95] Solter, K. M. & Gibor, A. Evidence for role of flagella as sensory transducers in mating of chlamydomonas reinhardi. Nature 265, 444–445 (1977).

[96] Taboada, C. et al. Glassfrogs conceal blood in their liver to maintain transparency. Science 378, 1315–1320 (2022).

[97] Schramma, N., Canales, G. C. & Jalaal, M. Light-regulated chloroplast morphodynamics in a single-celled dinoflagellate. Proc. Natl. Acad. Sci. U. S. A. 121, e2411725121 (2024).

[98] Stringer, C., Wang, T., Michaelos, M. & Pachitariu, M. Cellpose: a generalist algorithm for cellular segmentation. Nat. Methods 18, 100–106 (2021).

[99] Ershov, D. et al. TrackMate 7: integrating state-of-the-art segmentation algorithms into tracking pipelines. Nat. Methods 19, 829–832 (2022).

[100] Li, K. The image stabilizer plugin for imagej. https://www.cs.cmu.edu/~kangli/code/ImageStabilizer.html (2008).

[101] Gallet, B., Moriscot, C., Schoehn, G. & Decelle, J. Cryo-fixation and resin embedding of biological samples for electron microscopy and chemical imaging. protocols.io (2024).

