## Supplementary Information for "Long-range actin-driven endosymbiont mobility in a deep-diverging bilaterian"

Below are Supplementary Figures 1-6, which provide extensions of ideas presented in the main text.

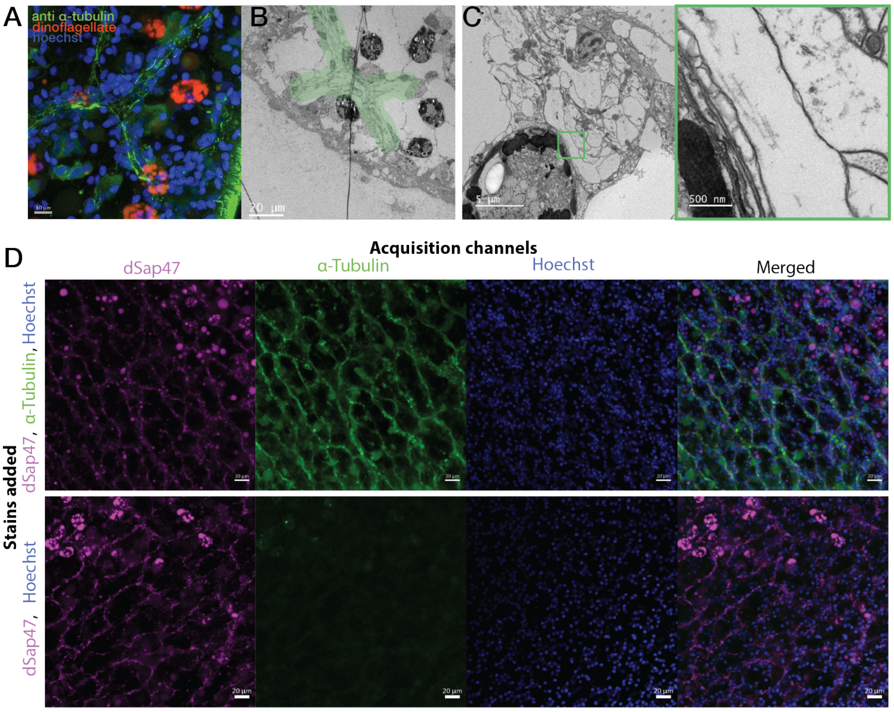


**Fig. 1** Polygonal cells that appear to guide direction of dinoflagellate mobility in ventral layer are host neurons (A) Anti-$\alpha$ tubulin staining reveals heavy tubulin content of polygonal cells visible in Fig.2D and SI Video 3 in the ventral layer. (B) TEM imaging on a similar area of a worm confirms neuron-like appearance of polygonal cells (green). (C) Higher magnification TEM imaging confirms that filaments in these cells are microtubules. (D) dSAP47 staining further confirms overlap between synaptotagmin stain and Anti-$\alpha$ tubulin, supporting the hypothesis that these polygonal shaped cells are host neurons. Control on the bottom row confirms that the Anti-$\alpha$ tubulin signal is not bleedthrough from dSAP47 signal.

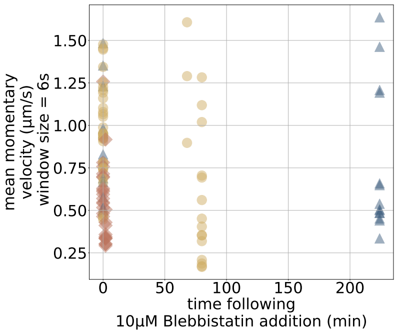


**Fig. 2** Addition of blebbistatin (Myosin II inhibitor) has no impact on symbiont mobility.

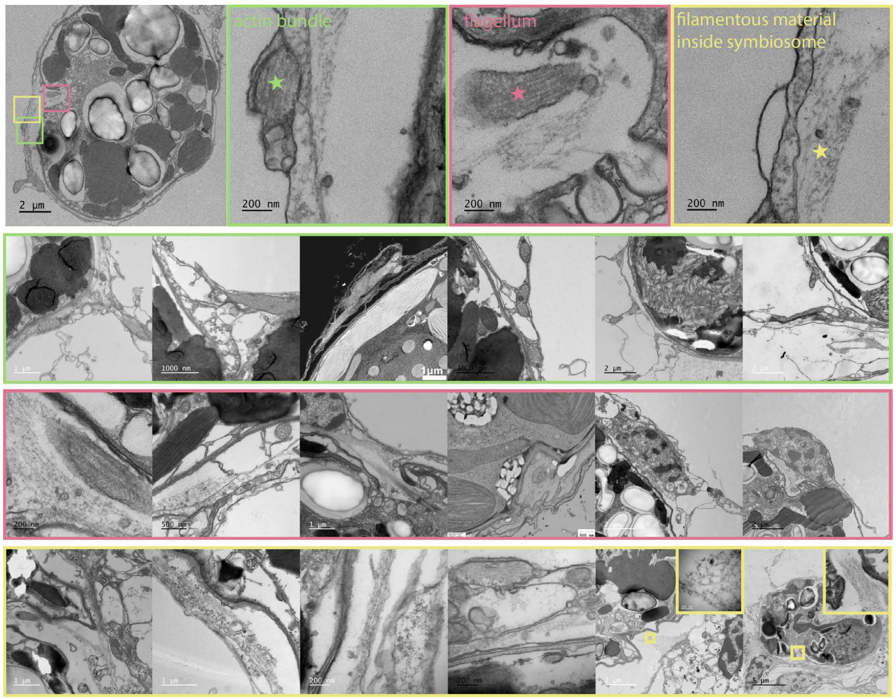


**Fig. 3** Additional details from TEM images on features discussed in main text. Unfortunately unbundled filamentous actin in cell cytoplasm that is hypothesized to be the main driver of symbiont mobility is difficult to see with traditional EM techniques. Boxed in green are examples in EM of bundled actin, which are part of host musculature (arrowheads in Main text Fig.4E). These are often found near symbionts because they penetrate the same negative spaces between large vacuolated central parenchyma cells but are not responsible for symbiont mobility. Boxed in pink are various examples of flagella seen in EM. In these images we can also see that host nuclei sometimes occupy the same negative spaces as symbionts. We also note an unidentified filmamentous material often found near flagella, inside the symbiosome. Boxed in yellow are more examples in EM of the filamentous material observed. Sometimes we see a clearly branched morphology (inset in bottom row, fifth image).

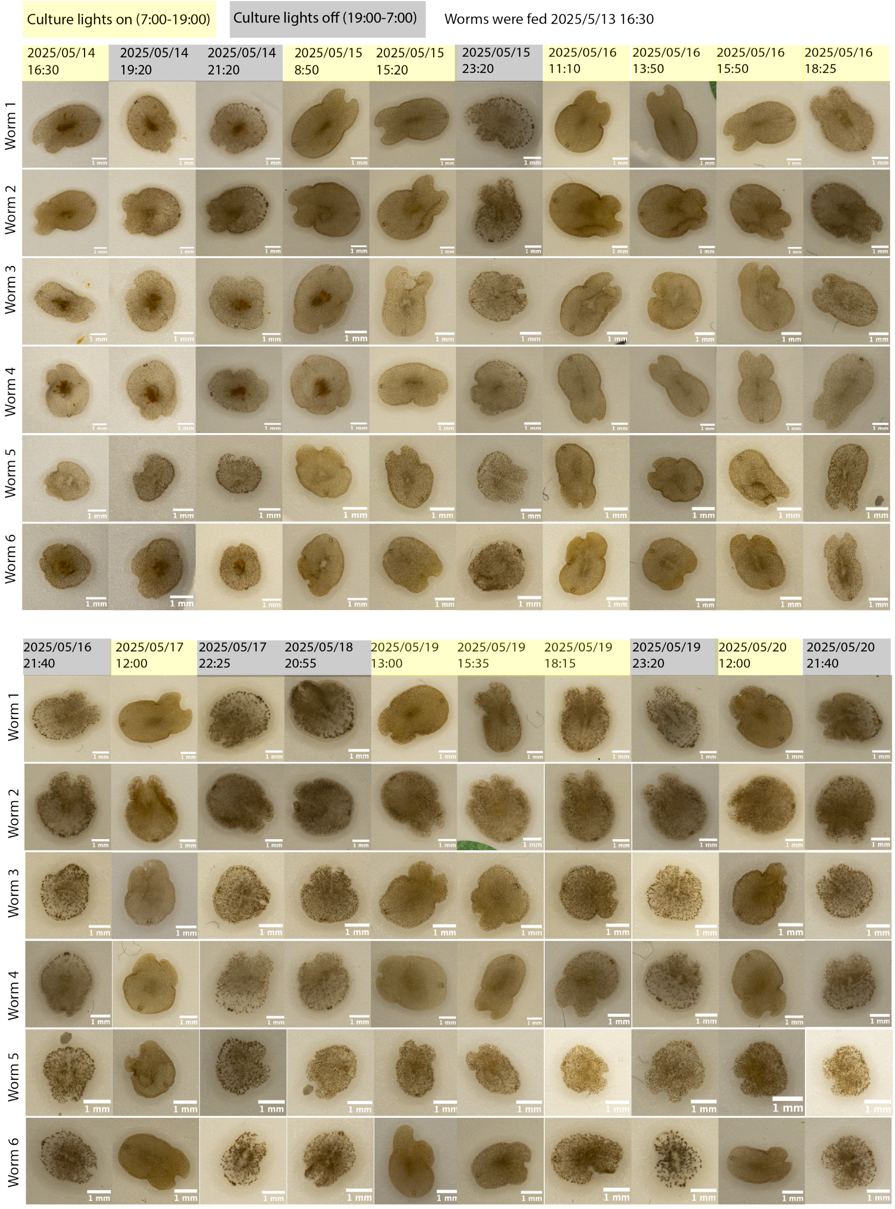


**Fig. 4** Macro-photography of six worms at various times of day confirms robustness of diel rhythms in symbiont distribution in whole, unamputated worms.

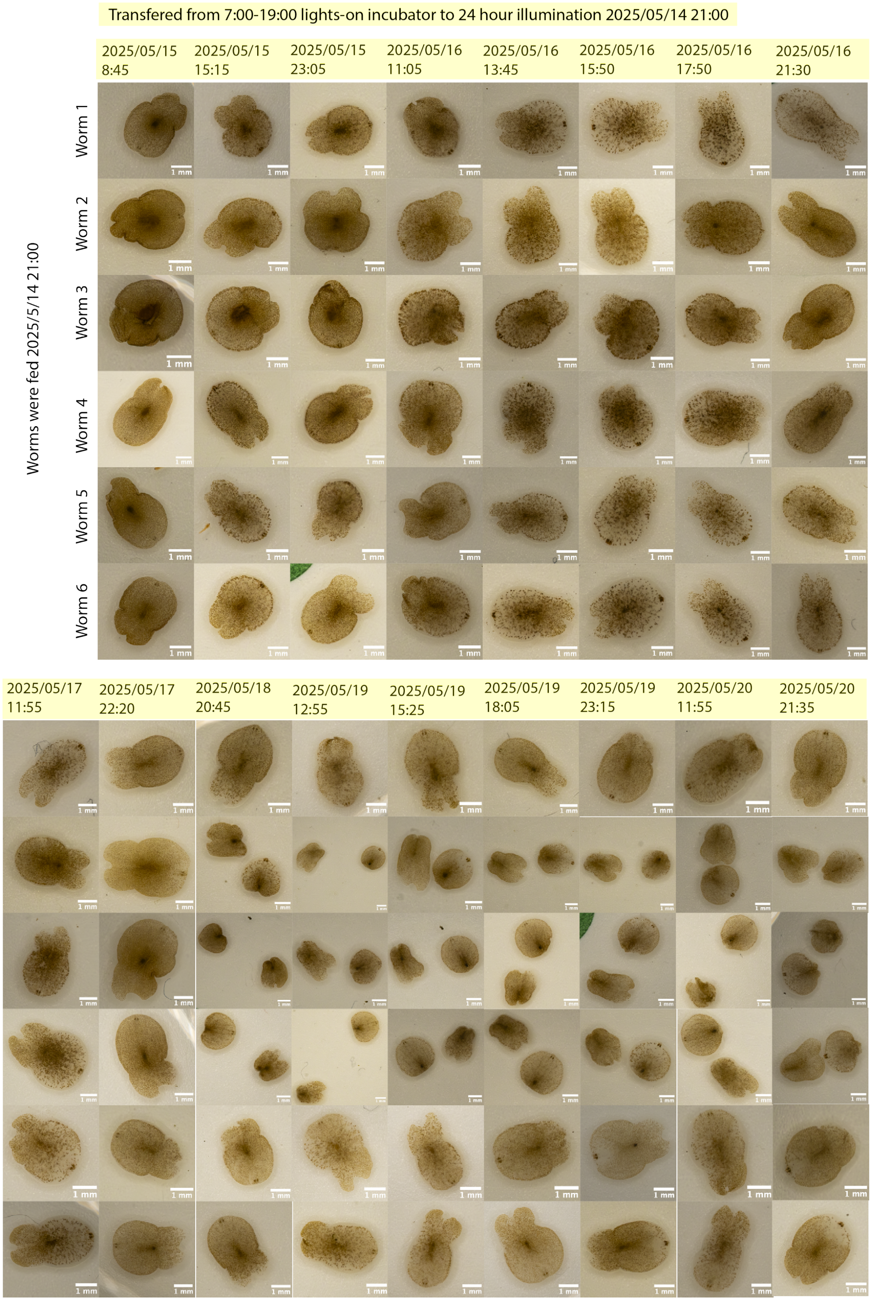


**Fig. 5** When worms are grown in 24-hour illumination, we still observe a diel rhythm in symbiont distribution, with a homogeneous distribution observed at some point during the day and clumping observed at other points. The rhythm appears disrupted - no longer cleanly homogenous during the day and clumping during the night - compared to worms grown in 12 hour - 12 hour light - dark cycle. This suggests other factors, such as light cycle, have an effect on diel rhythms observed.

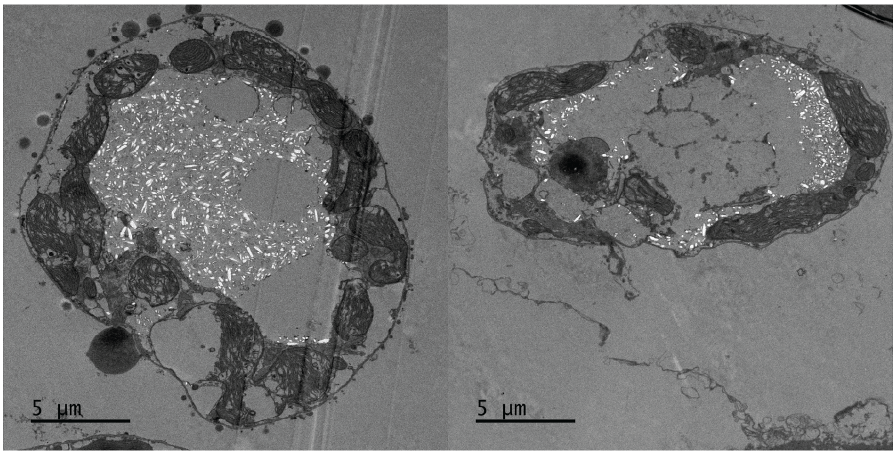


**Fig. 6** TEM imaging of worms cultured in the dark for two weeks shows potential symbiont digestion by host, as well as a clearly changed morphology of symbionts, with the presence of more crystal-like structures (perhaps starch).The absence of host membrane around symbionts may indicate that they have moved into the large vacuolated cells of the central parenchyma.
